# DNA-Encoded Multivalent Display of Protein Tetramers on Phage: Synthesis and *In Vivo* Aplications

**DOI:** 10.1101/2021.02.20.432100

**Authors:** Guilherme M. Lima, Alexey Atrazhev, Susmita Sarkar, Mirat Sojitra, Revathi Reddy, Matthew S. Macauley, Gisele Monteiro, Ratmir Derda

## Abstract

Phage display links phenotype of displayed polypeptides with DNA sequence in phage genome and offers a universal method for discovery of proteins with novel properties. Injection of phage-displayed libraries in living organisms further provides a unique and powerful approach to optimize biochemical, pharmacological and biological properties of the displayed peptides, antibodies and other proteins *in vivo*. However, over 60% of the proteome is comprised of multi-domain proteins, and display of large multi-subunit proteins on phages remains a challenge. Majority of protein display systems are based on monovalent phagemid constructs but methods for robust display of multiple copies of large proteins are scarce. Here, we describe a DNA-encoded display of a ∼200 kDa tetrameric protein tetrameric L-asparaginase on M13 phage produced by ligation of SpyCatcher-Asparaginase fusion (ScA) to prospectively barcoded phage clones displaying SpyTag peptide. Starting from the SpyTag display on p3 minor coat protein or p8 major coat protein yielded constructs with five copies of ScA displayed on p3 (ScA_5_-phage) and 50 copies of ScA on p8 protein (ScA_50_-phage). ScA remained active after conjugation. It could be easily produced directly from lysates of bacteria that express ScA. Display constructs of different valency can be injected into mice and analyzed by deep-sequencing of the DNA barcodes associated phage clones. In these multiplexed studies, we observed a density-dependent clearance rate *in vivo*. A known clearance mechanism of L-asparaginase is endocytosis by phagocytic cells. Our observations, thus, link the increase in density of the displayed protein with the increased rate of the endocytosis by cells *in vivo*. In conclusion, we demonstrate that a multivalent display of L-asparaginase on phage could be used to study the circulation life of this protein *in vivo* and such approach opens the possibility to use DNA sequencing to investigate multiplexed libraries of other multi-subunit proteins *in vivo*.

**Abstract Graphic:** 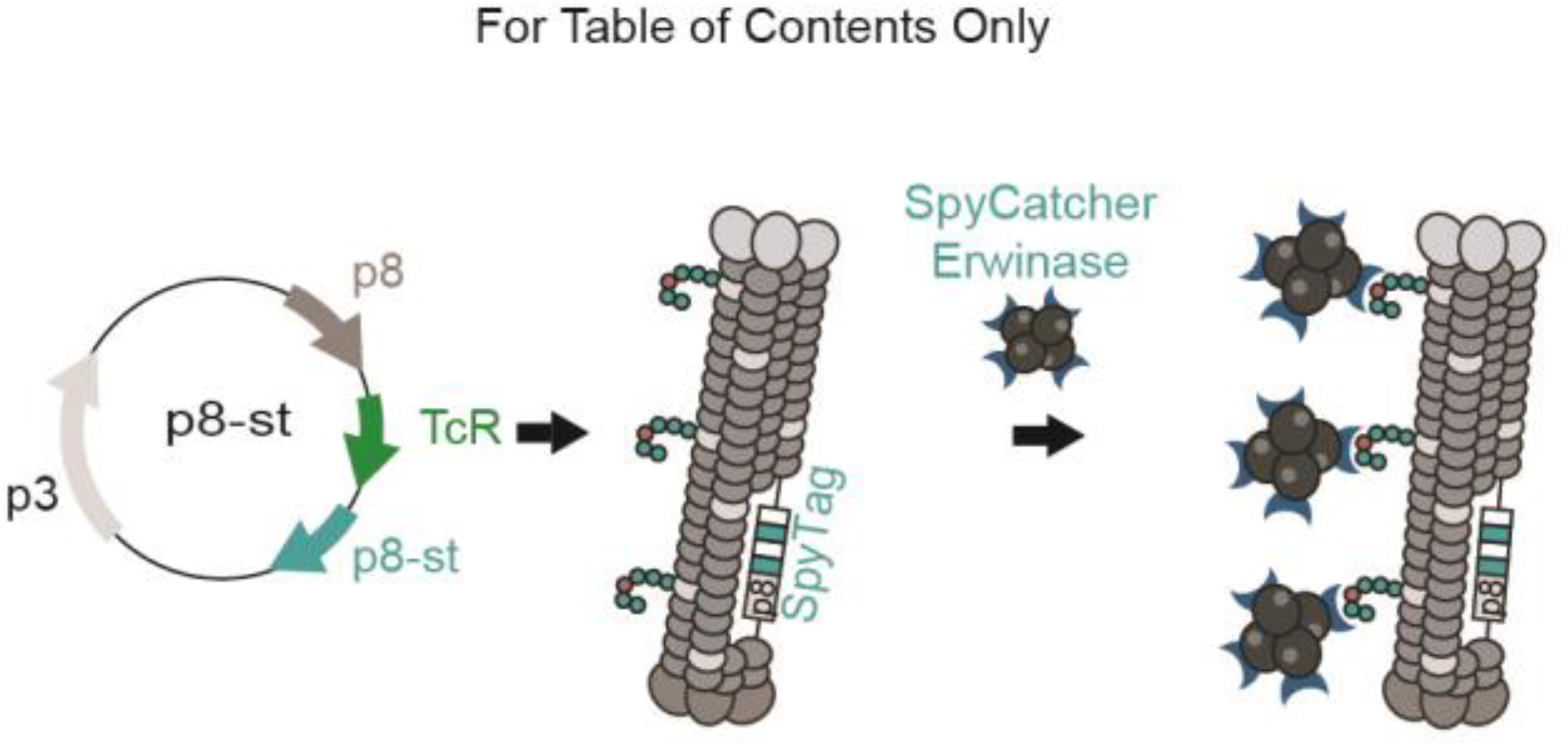

## Introduction

Display technologies such as yeast, phage and mRNA-display are standard tools for discovery of peptide and protein therapeutics^1^. Display technologies are poised for discovery and affinity maturation of ligands for purified protein targets and receptors on the surface of intact cell *in vitro*. Simple composition of M13 phage particles, their robustness to degradation and protection of DNA message by phage proteins makes it possible to use M13 phage display technology to discover ligands that bind to receptors *in vivo*. In multiple published reports, injection of M13-displyed libraries and DNA sequencing of phage clones recovered from specific organs led to identification of organ-homing peptides for liver,^2, 3^ heart, ^4, 5^ adipose tissue,^6^ brain,^7^ among others. Simultaneous identification of multiple pairs of targeted organs/peptides has also been accomplished using the same *in vivo* phage display approach.^8-10^ Such *in vivo* panning has been performed both in mice and in humans.^11^ Complementary *in vivo* panning can start from injection of phage in circulation and identify phage clones that do not home to any organs. Such approach led to discovery of peptide-decorated constructs that remain in circulation *in vivo* for a prolonged period of time.^12-14^ Both approaches suggest that phage display can be used to study and optimize *in vivo* pharmacokinetics of the peptide and protein therapeutic candidates. Molecules that are impossible to produce by standard translation machinery can also be displayed on phage.^15-17^ We recently demonstrated that DNA-encoded display of glycans on phage makes it possible to study the bio-distribution of glycans *in vivo*.^18^ In this manuscript, we build on this knowledge to produce multivalent, DNA-encoded display of multimeric proteins that cannot be expressed using traditional phage-display technology.

Over 60% of the proteome is comprised of multi-domain proteins; formation of homo-oligomers is critical for function of many proteins. For example, majority of glycan-binding proteins (lectins) exist as homo-dimers or higher order oligomers. To our knowledge only one example of phage display of a genetically encoded homo-tetrameric protein has been reported by Wells, Sidhu and co-workers.^19^ Methodology for display oligomeric proteins on phages would make it possible to use the benefits of phage display and *in vivo* display technologies to study these proteins. As the first demonstration, we focused on display of tetrameric enzyme L-asparaginase (ASNase). It is a 140 kDa C2-symmetric tetramer assembled from four identical monomers with four independent active sites that catalyse the hydrolysis of L-asparagine (L-Asn) to aspartic acid.^20^ ASNase is an important biopharmaceutical used in the treatment of acute lymphoblastic leukemia. Unlike normal blood cells, leukemic cells require extracellular L-Asn for protein synthesis because asparagine synthetase (ASNS) genes in these cells are epigenetically silenced.^21, 22^ Therefore, depletion of extracellular L-Asn by ASNase leads to inhibition of protein synthesis, starvation and selective apoptosis of cancer cells.^23-25^ Two forms of L-ASNase have been approved for the treatment of acute lymphoblastic leukemia: a native and PEGylated form from *Escherichia coli* and a native L-ASNase from *Erwinia chrysanthemi* (Erwinase). However, both non-PEGylated L-ASNase forms have short circulation lifetimes: 1.2 and 0.6 days, respectively.^26^ Furthermore, side effects of bacterial L-ASNases include neurotoxicity, allergic reactions that can lead to anaphylactic shock, and production of anti-asparaginase antibodies.^23, 27-30^ Development of new variants of ASNase can benefit from technologies that can encode and track a large number of variants of ASNase tetramers *in vivo*.

A plausible strategy for production of DNA-encoded display of protein tetramers builds on previously-reported conjugation of molecules to prospectively-barcoded bacteriophages and phage-displayed libraries of peptides. Enzymatic conjugation has been previously employed to conjugate proteins to M13 bacteriophage using Sortase^31^ or the SpyCatcher-SpyTag system.^32^ The latter SpyCatcher-SpyTag technology is a powerful approach for covalent conjugation of a 13 amino acid peptide termed SpyTag to a small 12.3 kDa protein domain termed SpyCatcher,^33-35^ with rates approaching the diffusion limit.^36^ Reaction successfully occurs within minutes and involves a nucleophilic attack of the Lys^31^ from SpyCatcher on the Asp^117^ from SpyTag, catalyzed by Glu^77^ in SpyCatcher.^35^ SpyTag and SpyCatcher can be incorporated both on C- and N-termini of the protein, whereas recognition by Sortase mandates display of the Sortase tag on the C-terminus of the protein. A sole report of SpyCatcher-SpyTag system in phage display employed a monovalent p3-display of SpyTag peptide or SpyCatcher domain in pFab5cHis phagemid vector to select SpyCatcher or SpyTag variants with increased reactivity.^32^ In this manuscript, we repurpose SpyTag and SpyCatcher technology to produce a multivalent DNA-encoded phage-display of proteins that cannot be produced using traditional display machinery. Starting from SpyCatcher-Erwinase fusion (ScA) and silently-encoded SpyTag-phage we displayed multiple copies of Erwinase tetramers on phage and evaluated the circulation time of these display constructs in mice. In the future, this technology can be employed to study the pharmacokinetics of DNA-encoded mixtures of Erwinase variants. It can be also expanded to display other multimeric proteins, such as lectins and study their bio-distribution *in vivo*.

## Results

### Conjugation of Erwinase to SpyTag-Displaying Phages

We successfully designed two phage constructs that display silently encoded SpyTag peptide libraries (Figure 1a, b and Supplementary Figure S1). The M13KE vector from New England Biolabs was used to display ∼5 copies of SpyTag peptide AHIVMVDAYKPTK,^35^ on minor protein p3 (p3-st_5_) (Supplementary Figure S1a). Type 88 phage-display vector fth1 developed by Gershoni group^37^ was used to display ∼50 copies of SpyTag on p8 (p8-st_50_) (Supplementary Figure S1b). Cloning of degenerated oligos coding for the peptide sequence AHIVMVDAYKPTK in each of the phage vectors yielded a library of up to 393,216 genotypically distinct phage clones of identical chemical composition (Figure 1a). This “silent barcoding”^17^ would allow tracking of the respective conjugate through phage sequencing, as previously done with barcoded glycan-decorated phages.^17, 18^ Individual phage clones were randomly picked and presence of the SpyTag sequence was confirmed by sequencing p3-st phage clones and p8-st-transformed bacterial colonies (Supplementary Figure S1). To show that peptide AHIVMVDAYKPTK does not interfere with export and incorporation of p8 into the extruding bacteriophage we used MALDI-TOF mass spectrometry to quantify the copy number of displayed peptides. MALDI confirmed a display of approximately 50 copies of the SpyTag peptide by the p8-st phage, which is a 10-fold increase in display density compared to the 5 copies in p3 display (Supplementary Figure S2).

**Figure 1.**
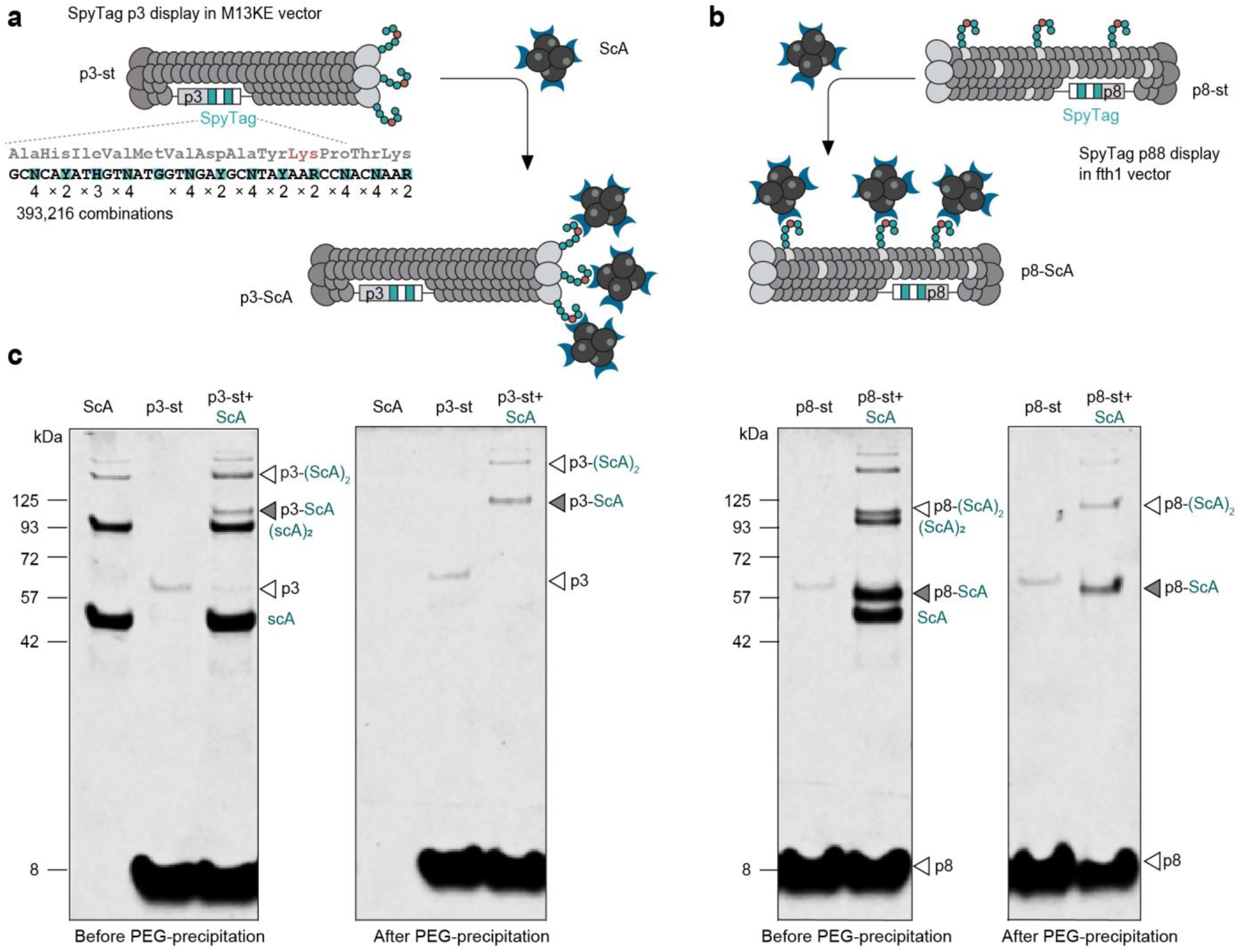
Conjugation of ScA to SpyTag-Expressing Phages. (a) SpyTag p3 display in M13KE vector (b) SpyTag p88 display in fth1 vector (c) SDS-PAGE image of conjugation reactions between ScA and p3-st phages (left-side images) or ScA and p8-st phages (right-side images). Grey arrows indicate protein-phage conjugates.

To produce a phage-display of a tetrameric Erwinase enzyme, we genetically fused the SpyCatcher domain to the N-terminus position of this protein and expressed SpyCatcher-Erwinase fusion protein (ScA) in *E. coli* as a soluble and enzymatically active protein (Supplementary Figure S3). SDS-PAGE gel analysis confirmed that p3-st_5_ covalently captures ScA (Figure 1c, grey arrows). Similarly, conjugation to the p8-st_50_ phage resulted in the formation of an ∼58 kDa band in SDS-PAGE gel (i.e., the sum of the molecular weight of the ScA monomer, p8 protein and the SpyTag peptide). These two new bands were not present when a control SVEKNDQKTYHA peptide (“SVEK”) was displayed on phage in place of SpyTag (Supplementary Figure S4). Ligation of ScA to p3-st consumed virtually all the available copies of the p3 protein, as confirmed by a drop in the intensity of the p3 band. This drop was not observed in SDS-PAGE of p8-display construct because recombinant p8-st band on SDS-PAGE gel was indistinguishable from the wild-type p8 protein band. However, MALDI analysis of p8-st_50_ phage before and after reaction with ScA-phage confirmed that the majority of the SpyTag-p8 peptides were consumed in the reaction. These results provided the first evidence for the production of p3- and p8-display constructs of a complex tetrameric enzyme. We refer to the constructs as p3-ScA_5_ and p8-ScA_50_ and employed several assays to confirm the display density.

Enzymatic properties of the Asparaginase allow convenient characterization of the display constructs. We purified p3-ScA_5_ and p8-ScA_50_ from the excess of unreacted ScA using PEG-NaCl precipitation,^31^ and confirmed the removal of the free ScA by SDS-PAGE (Figure 1c). The commonly used colorimetric Nessler assay^38^ demonstrated that both p3- and p8-constructs were enzymatically active (Figure 2a). Mixing the ScA sample with a phage that display the random SVEK peptide sequence or precipitation of p3-st and p8-st with no ScA presented, resulted in samples with no enzymatic activity (Figure 2a). We used Nessler assay and SDS-PAGE to demonstrate that p8-ScA can be produced simply by adding p8-st_50_ phage to a crude lysate of bacteria expressing ScA and purified by PEG-NaCl precipitation (Figure 3a). This approach successfully yielded the desired conjugates of reasonable purity: the characteristic ∼58 kDa p8-ScA conjugate band was observed. Other *E. coli* proteins originally present in the cell lysate were removed by PEG-NaCl purification (Figure 3b). The purified conjugate showed an enzymatic activity of 5.7 ± 0.8 U/mL, which is 44.4% of the original activity of the lysate (12.7 ± 1.6 U/mL) (Figure 3c). Taken together, these results demonstrate that it is possible to utilize ScA obtained directly from bacterial crude lysates with no column-based purification and a significant fraction of ScA proteins can be captured from the lysate.

**Figure 2.**
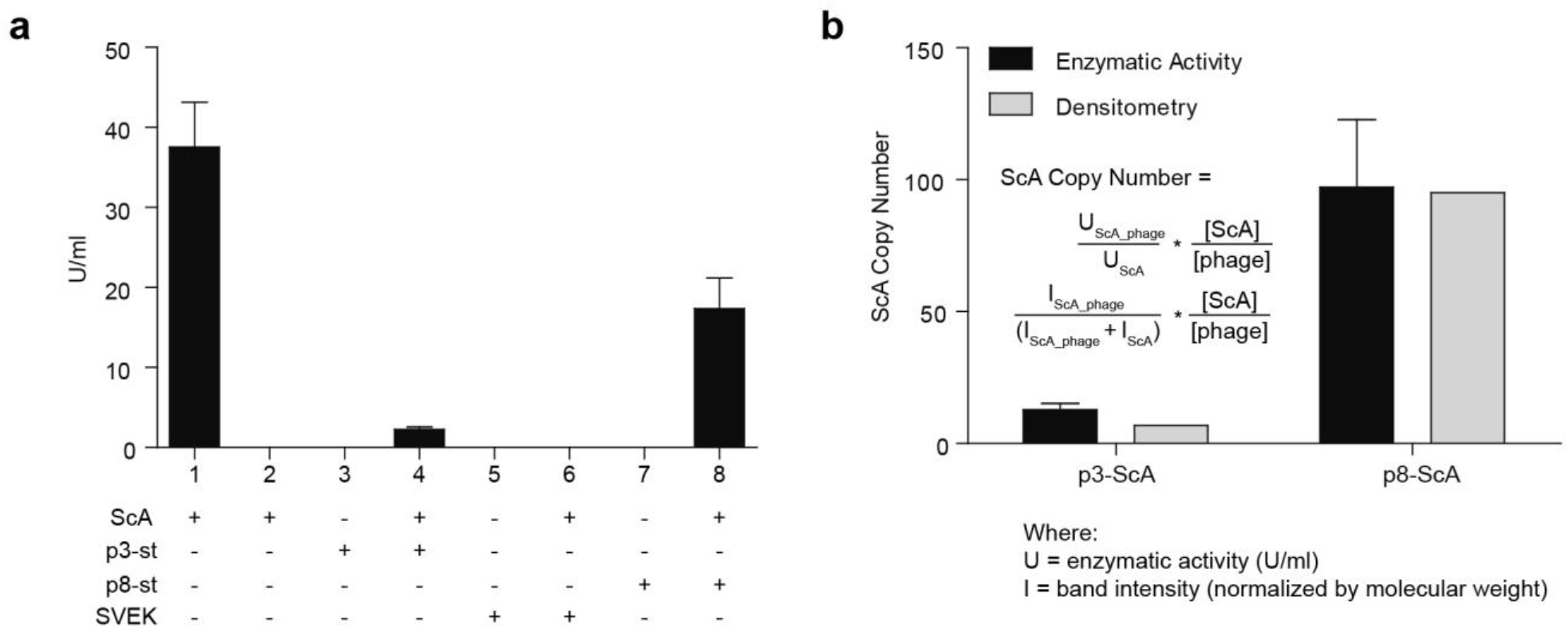
Characterization of ScA-phage conjugates obtained from purified ScA. (a) ScA enzymatic activity of purified ScA, purified ScA-phage conjugates and control conjugates. All reactions, except ‘1’, were precipitated using PEG/NaCl and pellet was later redissolved in PBS buffer. Activity of p3-ScA_5_ was reliably detected as 2.3 ± 0.3, whereas no detectable activity was observed for any controls (data is average ± standard deviation (std.), n=3) (d) Comparison of amount of protein-phage conjugate produced calculated either by enzymatic activity or SDS-PAGE image densitometry (data is average ± std. propagated from the std. of the enzymatic measurements n=3; n=1 is for densitometry).

**Figure 3.**
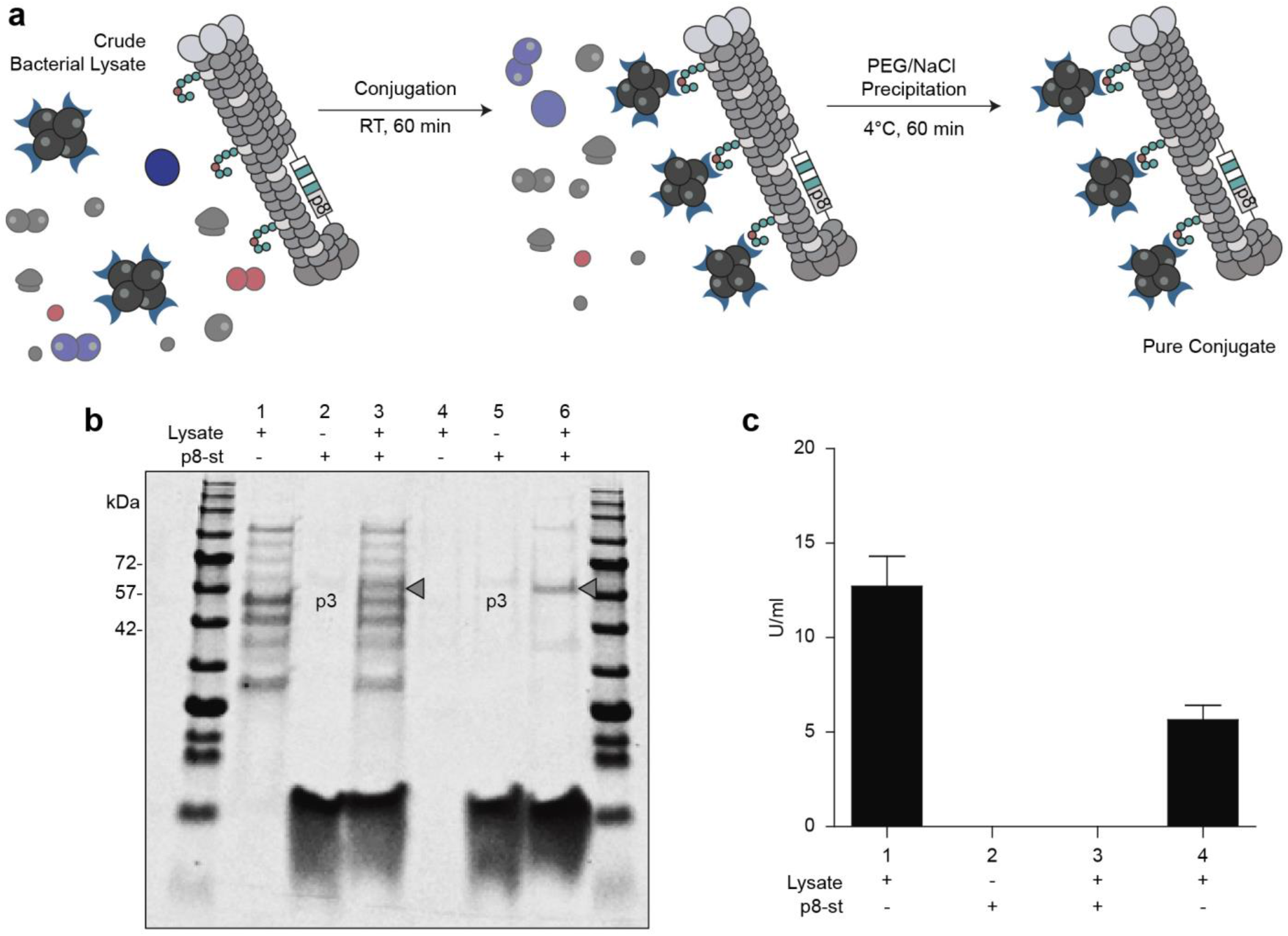
Characterization of ScA-phage conjugates obtained from crude lysate. (a) Scheme of protein-phage conjugate purification by PEG/NaCl precipitation. In this experiment, ScA-expressing crude bacterial lysates were mixed with p8-st phages to produce pure p8-ScA conjugates. (b) SDS-PAGE image showing conjugation reactions using a ScA-expressing lysate, before (Lanes 1, 2, and 3) and after purification by PEG/NaCl precipitation (Lanes 4, 5 and 6). Grey arrows indicate conjugates observed either before or after purification. (c) ScA enzymatic activity using ScA-expressing lysates. All reactions, except ‘1’, were precipitated using PEG/NaCl and pellet was later redissolved in PBS buffer. (data is average ± std.; n=3 measurements in independent enzymatic reactions).

### Characterization of the copy number of Erwinase on Phage

Accurate quantification of the copy number of active proteins displayed on phage is important for downstream applications. The quantification of enzymatic activity conveniently estimated the copy number of ScA molecules displayed on phage. From 17 ± 3.8 U/mL of enzymatic activity measured for samples containing p8-ScA_50_ and activity of the ScA alone, we extrapolated the copy number of enzymatically active ScA displayed in each these constructs (Figure 2a). Specifically, reaction of 3.3 nM of phage with 700 nM of ScA with 38 ± 5.6 U/mL enzymatic activity produced phage with 17 ± 3.8 U/mL enzymatic activity. The ratio of the activities multiplied by the original concentration of ScA yielded 320 ± 86 nM concentration of ScA associated with phage. Division of this number by phage concentration yielding (320 ± 86) nM / 3.3 nM = 97 ± 26 estimated copies of active ScA in p8-display. Recall that MALDI estimated approximately 50 copies of SpyTag in the p8-st construct (Supplementary Figure S2). A close agreement between MALDI (∼50 copies) and enzymatic activity (97 ± 26 copies) suggested that between 50 to 100 copies of ScA is displayed on phage. Repeating similar analysis for the p3-st construct that display up to 5 copies yielded a ScA copy number of 13 ± 2.4 (by enzymatic activity), 7 (by densitometry) (Figure 2b). The numbers were in agreement with the observation that 5 available copies of p3^39^ were consumed by ScA (Figure 1c).

A minor systematic increase observed in enzymatic assays may result from the presence of dimers of ScA. Minor amounts of dimers were observed during purification of ScA (Supplementary Figure S3) as well in the p3-ScA and p8-ScA phage conjugates (Figure 1c). This problem was caused by known self-reactivity of SpyCatcher domains^32^ (Supplementary Figure S5) and can be readily solved by using evolved versions of SpyCatcher/SpyTag.^36^ Despite minor presence of ScA dimers, the produced p3-ScA_5_ and p8-ScA_50_ construct to can be used to evaluate the *in vivo* clearance of Erwinase and the role of display valency in this clearance.

### In vivo Evaluation of ScA-phage Conjugates

The half-life of clearance of Erwinase in mice is ∼12 hours and receptor-mediated internalization by phagocytic cells, such as CD-14^+^ cells, is a known pathway for clearance of Erwinase.^40^ We hypothesized that increasing the density of the multivalent display leads to more effective engagement of these receptors and result in an apparent increase in the rate of clearance. To test this hypothesis, we evaluated *in vivo* clearance of the p3-ScA_5_ and p8-ScA_50_ conjugates of Erwinase (Figure 4a) and the role of protein density on phage particles on this clearance.

**Figure 4.**
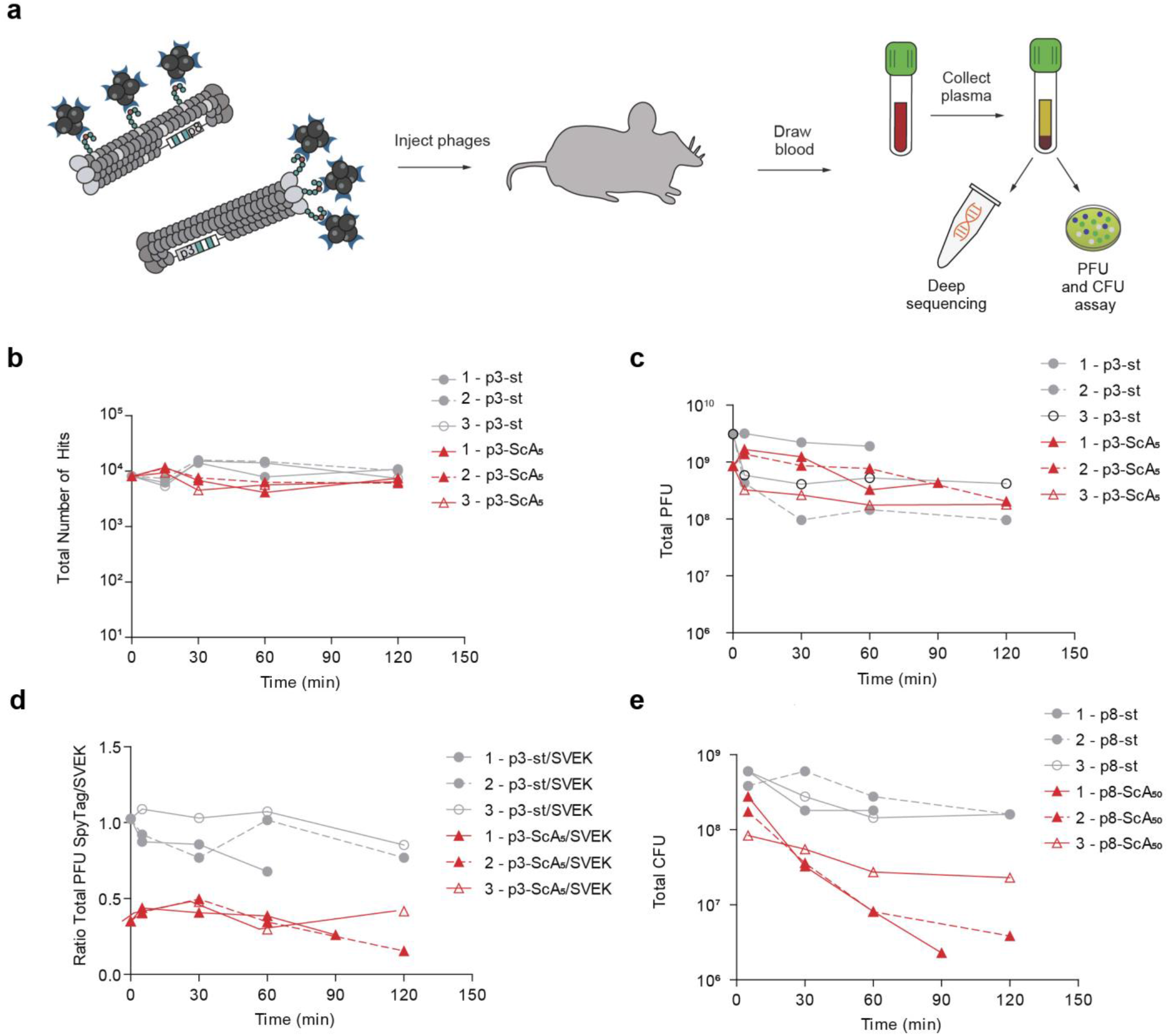
Clearance of protein-phage conjugates as measured by PFU and CFU. (a) Scheme of *in vivo* experiments. Protein-phage conjugates were injected into mice via the tail vein and blood samples were drawn at specific time points. Plasma samples were collected by centrifugation and then they were used for deep sequencing and titering analysis. (b) Clearance of p3-ScA_5_ phages from plasma as measured by the number of hits from each clone collected from deep sequencing data. (c) Clearance of p3-ScA_5_ phages from plasma as measured by counting the number of plaques in IPTG/X-Gal plates. (d) Clearance of p3-ScA_5_ phages from plasma as measured by the ratio of number of p3-st/p3-ScA_5_ plaques over the number of SVEK control plaques (p3-st/SVEK or p3-ScA_5_-/SVEK). Two phages were measured independently on the same agar overlay plate because they form plaques of two different color: p3-st and p3-ScA phages transduces LacZα and forms blue plaques in IPTG/X-Gal plates whereas SVEK transduces *GFP* gene and forms green-fluorescent plaques. Repeated Measures ANOVA statistical analysis was used to test for the influence of time and concentration differences between p3-ScA_5_ and p3-st phages from time = 0 (initial sample) to time = 60 min. (e) Clearance of p8-ScA_50_ with from plasma and control p8-st with no protein fused. p8-st and p8-ScA phage colonies were measured by the number of tetracycline-resistant colonies. Some colony counts were approximate. Colony count for samples 1 -p8-st (5 min), 3 – p8-st (5min) and 2 – p8-st (30 min) were > 5000 (Supplementary Table S2). Repeated Measures ANOVA statistical analysis was used to test for significance from time = 5 min to time = 60 min. Measurements are from three separate mice for each group. Average ratios of total PFU or total CFU were also plotted (Supplementary Figure S7).

We injected a mixture containing the p3-ScA_5_ (5×10^11^ PFU) conjugate and the SVEK phage (7×10^11^ PFU) in the tail vein of mice and measured the concentration of each phage clone in the blood by deep sequencing. Phage clone not conjugated to ScA (p3-st, 6×10^11^ PFU) was used as a control. Once plasma samples were diluted (1:1000), the downstream PCR and quantification of DNA were not influenced by the presence of residual plasma components (Supplementary Figure S6b). PCR amplification of phage genome from the collected plasma samples without any extensive manipulation is important as purification or amplification of phage may introduce undesired biases. Deep sequencing demonstrated no significant differences between the copy number of p3-ScA_5_ and control p3-st samples (Figure 4b). This observation suggested that presence of 5 copies of ScA on phage does not lead to substantial increase clearance of phage. Titering st-p3, p3-ScA_5_ and SVEK phages in plasma corroborated these results (Figure 4c). SVEK and st-p3 phage were engineered to form plaques of different color^18^ and allow simultaneous quantification in the same mixture. Normalization by the titer of SVEK phage present in each injection dampened the mouse-to-mouse biological variability and again, confirmed that the presence of SpyTag or ScA on phage does not lead to significant differences in clearance of phage (Figure 4d).

In contrast to p3-ScA_5_, p8-ScA_50_ phage constructs exhibited a significant increase in clearance when compared to phage that contained only a p8-displayed SpyTag (p8-st_50_) (Figure 4e). The concentration of p8-ScA_50_ consistently decreased in each of the three independent serum samples collected from three treated mice. Total p8-ScA_50_ concentration started at around 10^8^ CFU (colony forming units) and dropped down to approximately 10^6^-10^7^ CFU. On the other hand, the concentration of p8-st_50_ phage without ScA exhibited only a minor decrease during the 120-min of the *in vivo* experiment and remained above 10^8^ CFU. These data demonstrate that phage particles that displayed a significant higher copy number of ScA—50 copies for p8-ScA-p8 as opposed to 5 for p3-ScA—were cleared faster from circulation (p<0.01). Although we did not examine the biodistribution of p8-ScA_50_ in this manuscript, the observation is consistent with the proposal that higher density of the multivalent display should lead to an increased internalization by phagocytic cells and lead to an apparent in increase in the rate of clearance. Future experiments can employ deep-sequencing of phage isolated from CD-14^+^ cells, other immune cells as well as organs (e.g., liver) to expand on this proposal.

## Discussion

Binding of multivalent ligands to cellular receptors controls many biological mechanisms.^41^ In this study, an efficiently display of multiple copies of Erwinase on the surface of filamentous phages, demonstrated investigation of the effect of density of display on the interaction of the displayed proteins with cellular receptors *in vivo*. One of most important clearance mechanisms of ASNase is endocytosis by phagocytic cells.^40^ ASNase molecules rapidly accumulate in macrophage-rich tissues where they are internalized by these cells and degraded by the lysosomal protease cathepsin B.^40, 42^ This study has demonstrated that high-copy number p8-display of Erwinase can dramatically increase the clearance of the corresponding protein-phage conjugate in comparison to the low-copy number p3-display of Erwinase. This result is in agreement with previous studies of Finn, Baird and Kiessling groups in which polyvalent display of biomolecules on viral particles can potentiate receptor-mediated cellular uptake.^43, 44^

SpyCatcher-SpyTag system^35^ effectively ligated ∼200 kDa tetrameric enzyme to the N-terminus position of both the p3 and the p8 viral capsid proteins. Despite the relatively large molecular weight of ScA, our biological conjugation approach allowed us to display 50 copies of this protein on phage, with no significant impact on the biological activity of the displayed enzyme. Two other examples of display of multi-domain proteins on phage exist.^19, 31^ Guimaraes and coworkers used sortase-mediated reactions to functionalize p3 with recombinant streptavidin, a homo-tetrameric protein achieving display of ∼5 copies.^31^ Sidhu, Weiss and Wells displayed streptavidin tetramer on p8 by genetically fusing one of streptavidin subunits to the N-terminus of a mutated p8 protein.^19^ The authors used phage ELISA to elegantly demonstrated that specific mutations in p8 increase the relative display density on p8 (the authors re-optimized the p8 sequence for every class of displayed protein). However, the manuscript provided no information about the absolute number of the displayed streptavidin molecules. Traditional phage-display approach mandates expression and periplasmic export of displayed proteins in the same bacteria cell as the phage itself. The SpyCatcher-SpyTag framework allows production of high copy number phage-display of proteins that originate from any source. The expression of the fusion of SpyTag and protein of interest can be performed in a different organism making it possible to access a larger diversity of protein structures as well as proteins with glycosylation and other post-translational modifications not accessible in *E. coli*. As of today, dozens of articles exploiting the use of SpyCatcher fusion proteins have been published in the literature^45-47^; all of these proteins should be applicable to display on M13-SpyTag particle.

Conjugation reaction occurred efficiently at high nanomolar concentrations (700 nM) of purified ScA or even lower concentrations of ScA present on the crude bacterial lysate (240 nM of non-purified ScA). The latter calculation was based on enzymatic activity of ScA in the crude lysate relative to the enzymatic activity of purified ScA (Figures 2a, 3c). Recently, Howarth group reported a new, more sensitive version of SpyCatcher and SpyTag pairs that can react with each other at low nanomolar concentrations: a significant increase in reaction rate compared to the system currently used in this work.^36^ Other versions of SpyCatcher^32^ also minimize the formation of undesirable oligomers observed in our system. One of the potential problems in this report, is the need to introduce a ∼12 kDa protein domain in the tetrameric protein. Designing phage that can display SpyCatcher rather than SpyTag would eliminate this problem, as the protein partner would have to be fused to the short 13-mer SpyTag peptide rather than a ∼12kDa protein domain. This problem could also be solved by implementing the SnoopCatcher/SnoopTag reactive pairs.^48^

It was interesting that the display of large tetrameric proteins did not ablate the infectivity of phage but resulted only in minor decrease in infectivity. The ratio of PFU from p3-st/SVEK and p3-ScA_5_ /SVEK was ∼2-fold lower for phages containing conjugated ScA than non-conjugated phages (p<0.05) (Figure 4d); whereas deep-sequencing and qPCR results suggested a similar copy number of phage DNA. This observation might be explained by a minor, 2-fold loss of phage infectivity caused by attachment of ScA to the p3 protein. The p3 protein is responsible for phage attachment to the bacterial pilus during infection^39^ and ScA might be sterically hindering the interaction between p3 and the bacterial pilus.^49^ Diversification of phage “post-expression” with proteins of various sizes can be used to revisit this hypothesis and decouple phage infectivity from phage expression in a manner that isn’t accessible to traditional phage display systems.

Display of multiple protein copies on viral or virus-like particles is a growing field due to its promise in areas such as protein-based vaccines.^14, 50-52^ A recent example includes immunization of mice with multiple SARS-CoV-2 spike glycoprotein antigens conjugated to a virus-like particle using the SpyCatcher-SpyTag system.^53^ Using the genetically barcoded phage in such systems as a tool may offer a significant benefit. It allows production of various conjugates that are tested simultaneously as one mixture of multiple conjugates *in vivo*. Unlike traditional genetic encoding that produces billion-scale libraries, silent barcoding is suitable to produce libraries of ∼100 variants.^18^ Benefits of 100x-mutiplexing is significant nevertheless. Injection of the “cassette” of ∼100 barcoded conjugates *in vivo* can be used to decrease the number of animals required for experiments and introduce critical internal controls. *In vivo* injection of multivalent constructs that display glycan-binding proteins (lectins) would be a powerful tool to study *in vivo* glycobiology; such approach has not yet been exploited yet but closely related DNA-encoded lectin array^54^ or phage-displayed glycan array^18^ technologies have been reported recently. Many natural lectins function only as homodimers, trimers or tetramers. DNA-encoded multivalent presentation of such lectins could be readily built on the methodology described in this paper. Interactions of glycans with lectins occurs with dissociation constants of low millimolar and high micromolar ranges^55^, and it is enhanced by multivalent presentation of these lectins.^41, 56-63^ DNA-barcoded multivalent presentation of libraries of lectins on phage—akin to DNA-barcoded “liquid glycan array”^18^—would make it possible to profile weak interactions between GBPs and glycans on the surface of cells *in vivo*. Our group is currently working in implementing such technology and results will be presented in subsequent manuscripts. The multiplexing capability of this technology can advance other potential biomedical applications which include the use of protein-bacteriophage conjugates as means to assess, *in vivo*, their biodistribution,^64^ cellular uptake^65^ and blood retention time.^14^

## Acknowledgments

We thank the staff at the University of Alberta mass spectrometry facility (Chemistry Department) for help with MALDI analysis and Sophie Dang at the molecular biology service unit for assistance with Illumina sequencing. The authors acknowledge funding from NSERC (RGPIN-2016-402511 to R.D. and RGPIN-2018-03815 to M.M.), Alberta Innovates Strategic Research Project to R.D., GlycoNet (CD-03 to M.M and R.D.), São Paulo Research Foundation (FAPESP, Brazil, process numbers: 2018/15041-8, 2019/09354-6, 2018/15104-0 and 2013/08617-7 to G.M) and a research productivity fellowship from CNPq 309224/2019-5 CAPES finance code 001 (to G.M.). Infrastructure support was provided by CFI New Leader Opportunity (to R.D). We thank Gareth Lambkin at the University of Alberta Biological Services for assistance with protein purification and provision of biological instruments. We also thank Dr. Jonathan M. Gersoni, from Tel Aviv University, Israel, for his gift of plasmid fth1.

## Supporting Information

Supplementary document contains Material and Methods, Supplementary Figures (S1-S8) and Supplementary Tables (S1-S2).

## References

1. Sergeeva, A., Kolonin, M. G., Molldrem, J. J., Pasqualini, R., Arap, W., Display technologies: application for the discovery of drug and gene delivery agents. Adv. Drug Del. Rev. 2006, 58 (15), 1622–1654.

2. Du, B., Han, H., Wang, Z., Kuang, L., Wang, L., Yu, L., Wu, M., Zhou, Z., Qian, M., Targeted drug delivery to hepatocarcinoma in vivo by phage-displayed specific binding peptide. Mol. Cancer Res. 2010, 8 (2), 135–44.

3. Ludtke, J. J., Sololoff, A. V., Wong, S. C., Zhang, G., Wolff, J. A., In vivo selection and validation of liver-specific ligands using a new T7 phage peptide display system. Drug Deliv. 2007, 14 (6), 357–69.

4. Zahid, M., Phillips, B. E., Albers, S. M., Giannoukakis, N., Watkins, S. C., Robbins, P. D., Identification of a cardiac specific protein transduction domain by in vivo biopanning using a M13 phage peptide display library in mice. PLoS One 2010, 5 (8), e12252.

5. Kanki, S., Jaalouk, D. E., Lee, S., Yu, A. Y., Gannon, J., Lee, R. T., Identification of targeting peptides for ischemic myocardium by in vivo phage display. J. Mol. Cell. Cardiol. 2011, 50 (5), 841–8.

6. Lee, N. K., Kim, H. S., Kim, K. H., Kim, E. B., Cho, C. S., Kang, S. K., Choi, Y. J., Identification of a novel peptide ligand targeting visceral adipose tissue via transdermal route by in vivo phage display. J. Drug Target. 2011, 19 (9), 805–13.

7. Li, J., Zhang, Q., Pang, Z., Wang, Y., Liu, Q., Guo, L., Jiang, X., Identification of peptide sequences that target to the brain using in vivo phage display. Amino Acids 2012, 42 (6), 2373–81.

8. Rajotte, D., Arap, W., Hagedorn, M., Koivunen, E., Pasqualini, R., Ruoslahti, E., Molecular heterogeneity of the vascular endothelium revealed by in vivo phage display. J. Clin. Invest. 1998, 102 (2), 430–7.

9. Kolonin, M. G., Sun, J., Do, K. A., Vidal, C. I., Ji, Y., Baggerly, K. A., Pasqualini, R., Arap, W., Synchronous selection of homing peptides for multiple tissues by in vivo phage display. FASEB J. 2006, 20 (7), 979–81.

10. Arap, W., Kolonin, M. G., Trepel, M., Lahdenranta, J., Cardó-Vila, M., Giordano, R. J., Mintz, P. J., Ardelt, P. U., Yao, V. J., Vidal, C. I., Chen, L., Flamm, A., Valtanen, H., Weavind, L. M., Hicks, M. E., Pollock, R. E., Botz, G. H., Bucana, C. D., Koivunen, E., Cahill, D., Troncoso, P., Baggerly, K. A., Pentz, R. D., Do, K.-A., Logothetis, C. J., Pasqualini, R., Steps toward mapping the human vasculature by phage display. Nat. Med. 2002, 8, 121–127.

11. Bábíčková, J., Tóthová, Ľ., Boor, P., Celec, P., In vivo phage display—a discovery tool in molecular biomedicine. Biotechnol. Adv. 2013, 31 (8), 1247–1259.

12. Merril, C. R., Biswas, B., Carlton, R., Jensen, N. C., Creed, G. J., Zullo, S., Adhya, S., Long-circulating bacteriophage as antibacterial agents. Proc. Natl. Acad. Sci. U. S. A. 1996, 93 (8), 3188–92.

13. Sokoloff, A. V., Bock, I., Zhang, G., Sebestyen, M. G., Wolff, J. A., The interactions of peptides with the innate immune system studied with use of T7 phage peptide display. Mol. Ther. 2000, 2 (2), 131–9.

14. Jin, P., Sha, R., Zhang, Y., Liu, L., Bian, Y., Qian, J., Qian, J., Lin, J., Ishimwe, N., Hu, Y., Zhang, W., Liu, Y., Yin, S., Ren, L., Wen, L. P., Blood Circulation-Prolonging Peptides for Engineered Nanoparticles Identified via Phage Display. Nano Lett. 2019, 19 (3), 1467–1478.

15. Vinals, D. F., Kitov, P. I., Tu, Z., Zou, C., Cairo, C. W., Lin, H. C.-H., Derda, R., Selection of galectin-3 ligands derived from genetically encoded glycopeptide libraries. Peptide Science 2019, 111 (1), e24097.

16. Ng, S., Bennett, N. J., Schulze, J., Gao, N., Rademacher, C., Derda, R., Genetically-encoded Fragment-based Discovery of Glycopeptide Ligands for DC-SIGN. Biorg. Med. Chem. 2018.

17. Tjhung, K. F., Kitov, P. I., Ng, S., Kitova, E. N., Deng, L., Klassen, J. S., Derda, R., Silent Encoding of Chemical Post-Translational Modifications in Phage-Displayed Libraries. Journal of the American Chemical Society 2016, 138 (1), 32-35.

18. Sojitra, M., Sarkar, S., Maghera, J., Rodrigues, E., Carpenter, E., Seth, S., Vinals, D. F., Bennett, N., Reddy, R., Khalil, A., Xue, X., Bell, M., Zheng, R. B., Zhang, P., Nycholat, C., Ling, C.-C., Lowary, T. L., Paulson, J. C., Macauley, M. S., Derda, R., Genetically Encoded, Multivalent Liquid Glycan Array (LiGA). bioRxiv 2020, 2020.03.24.997536.

19. Sidhu, S. S., Weiss, G. A., Wells, J. A., High copy display of large proteins on phage for functional selections. J. Mol. Biol. 2000, 296 (2), 487–95.

20. Aghaiypour, K., Wlodawer, A., Lubkowski, J., Structural basis for the activity and substrate specificity of Erwinia chrysanthemi L-asparaginase. Biochemistry 2001, 40 (19), 5655–64.

21. Akagi, T., Yin, D., Kawamata, N., Bartram, C. R., Hofmann, W. K., Wolf, I., Miller, C. W., Koeffler, H. P., Methylation analysis of asparagine synthetase gene in acute lymphoblastic leukemia cells. Leukemia 2006, 20 (7), 1303–6.

22. Pastorczak, A., Fendler, W., Zalewska-Szewczyk, B., Gorniak, P., Lejman, M., Trelinska, J., Walenciak, J., Kowalczyk, J., Szczepanski, T., Mlynarski, W., Polish Pediatric Leukemia/Lymphoma Study, G., Asparagine synthetase (ASNS) gene polymorphism is associated with the outcome of childhood acute lymphoblastic leukemia by affecting early response to treatment. Leuk. Res. 2014, 38 (2), 180–3.

23. Narta, U. K., Kanwar, S. S., Azmi, W., Pharmacological and clinical evaluation of L-asparaginase in the treatment of leukemia. Crit. Rev. Oncol. Hematol. 2007, 61 (3), 208–21.

24. Avramis, V. I., Asparaginases: biochemical pharmacology and modes of drug resistance. Anticancer Res. 2012, 32 (7), 2423–37.

25. Emadi, A., Zokaee, H., Sausville, E. A., Asparaginase in the treatment of non-ALL hematologic malignancies. Cancer Chemother. Pharmacol. 2014, 73 (5), 875–83.

26. Asselin, B. L., The three asparaginases. Comparative pharmacology and optimal use in childhood leukemia. Adv Exp Med Biol 1999, 457, 621–9.

27. Narazaki, H., Kaizu, K., Miyatake, C., Koizumi, S., Asano, T., Fujino, O., Delayed-type hypersensitivity in response to L-asparaginase in a case of acute lymphoblastic leukemia. J Nippon Med Sch 2012, 79 (6), 489–93.

28. Pieters, R., Hunger, S. P., Boos, J., Rizzari, C., Silverman, L., Baruchel, A., Goekbuget, N., Schrappe, M., Pui, C. H., L-asparaginase treatment in acute lymphoblastic leukemia: a focus on Erwinia asparaginase. Cancer 2011, 117 (2), 238–49.

29. Raetz, E. A., Salzer, W. L., Tolerability and efficacy of L-asparaginase therapy in pediatric patients with acute lymphoblastic leukemia. J. Pediatr. Hematol. Oncol. 2010, 32 (7), 554–63.

30. Rizzari, C., Conter, V., Stary, J., Colombini, A., Moericke, A., Schrappe, M., Optimizing asparaginase therapy for acute lymphoblastic leukemia. Curr. Opin. Oncol. 2013, 25 Suppl 1, S1–9.

31. Hess, G. T., Cragnolini, J. J., Popp, M. W., Allen, M. A., Dougan, S. K., Spooner, E., Ploegh, H. L., Belcher, A. M., Guimaraes, C. P., M13 bacteriophage display framework that allows sortase-mediated modification of surface-accessible phage proteins. Bioconjug. Chem. 2012, 23 (7), 1478–87.

32. Keeble, A. H., Banerjee, A., Ferla, M. P., Reddington, S. C., Anuar, I., Howarth, M., Evolving Accelerated Amidation by SpyTag/SpyCatcher to Analyze Membrane Dynamics. Angewandte Chemie-International Edition 2017, 56 (52), 16521–16525.

33. Liu, Y., Liu, D., Yang, W., Wu, X. L., Lai, L., Zhang, W. B., Tuning SpyTag-SpyCatcher mutant pairs toward orthogonal reactivity encryption. Chem. Sci. 2017, 8 (9), 6577–6582.

34. Zhang, W. B., Sun, F., Tirrell, D. A., Arnold, F. H., Controlling macromolecular topology with genetically encoded SpyTag-SpyCatcher chemistry. J. Am. Chem. Soc. 2013, 135 (37), 13988–97.

35. Zakeri, B., Fierer, J. O., Celik, E., Chittock, E. C., Schwarz-Linek, U., Moy, V. T., Howarth, M., Peptide tag forming a rapid covalent bond to a protein, through engineering a bacterial adhesin. Proc. Natl. Acad. Sci. U. S. A. 2012, 109 (12), E690–E697.

36. Keeble, A. H., Turkki, P., Stokes, S., Anuar, I., Rahikainen, R., Hytonen, V. P., Howarth, M., Approaching infinite affinity through engineering of peptide-protein interaction. Proc. Natl. Acad. Sci. U. S. A. 2019, 116 (52), 26523–26533.

37. Enshell-Seijffers, D., Smelyanski, L., Gershoni, J. M., The rational design of a ’type 88’ genetically stable peptide display vector in the filamentous bacteriophage fd. Nucleic Acids Res. 2001, 29 (10), E50–0.

38. Costa-Silva, T. A., Costa, I. M., Biasoto, H. P., Lima, G. M., Silva, C., Pessoa, A., Monteiro, G., Critical overview of the main features and techniques used for the evaluation of the clinical applicability of L-asparaginase as a biopharmaceutical to treat blood cancer. Blood Rev. 2020, 43.

39. Smith, G. P., Petrenko, V. A., Phage display. Chem. Rev. 1997, 97 (2), 391–410.

40. van der Meer, L. T., Terry, S. Y., van Ingen Schenau, D. S., Andree, K. C., Franssen, G. M., Roeleveld, D. M., Metselaar, J. M., Reinheckel, T., Hoogerbrugge, P. M., Boerman, O. C., van Leeuwen, F. N., In Vivo Imaging of Antileukemic Drug Asparaginase Reveals a Rapid Macrophage-Mediated Clearance from the Bone Marrow. J. Nucl. Med. 2017, 58 (2), 214–220.

41. Kiessling, L. L., Gestwicki, J. E., Strong, L. E., Synthetic multivalent ligands as probes of signal transduction. Angewandte Chemie-International Edition 2006, 45 (15), 2348–2368.

42. Van Der Meer, L. T., Waanders, E., Levers, M., Venselaar, H., Roeleveld, D., Boos, J., Lanvers, C., Brüggemann, R. J., Kuiper, R. P., Hoogerbrugge, P. M., A germ line mutation in cathepsin B points toward a role in asparaginase pharmacokinetics. Blood, The Journal of the American Society of Hematology 2014, 124 (19), 3027–3029.

43. Banerjee, D., Liu, A. P., Voss, N. R., Schmid, S. L., Finn, M. G., Multivalent display and receptor-mediated endocytosis of transferrin on virus-like particles. ChemBioChem 2010, 11 (9), 1273–9.

44. Larocca, D., Jensen-Pergakes, K., Burg, M. A., Baird, A., Receptor-targeted gene delivery using multivalent phagemid particles. Mol. Ther. 2001, 3 (4), 476–84.

45. Pardee, K., Slomovic, S., Nguyen, P. Q., Lee, J. W., Donghia, N., Burrill, D., Ferrante, T., McSorley, F. R., Furuta, Y., Vernet, A., Portable, on-demand biomolecular manufacturing. Cell 2016, 167 (1), 248-259. e12.

46. Sun, F., Zhang, W.-B., Mahdavi, A., Arnold, F. H., Tirrell, D. A., Synthesis of bioactive protein hydrogels by genetically encoded SpyTag-SpyCatcher chemistry. Proceedings of the National Academy of Sciences 2014, 111 (31), 11269–11274.

47. Nguyen, P. Q., Botyanszki, Z., Tay, P. K. R., Joshi, N. S., Programmable biofilm-based materials from engineered curli nanofibres. Nature communications 2014, 5 (1), 1–10.

48. Veggiani, G., Nakamura, T., Brenner, M. D., Gayet, R. V., Yan, J., Robinson, C. V., Howarth, M., Programmable polyproteams built using twin peptide superglues. Proc. Natl. Acad. Sci. U. S. A. 2016, 113 (5), 1202–1207.

49. Carmen, S., Jermutus, L., Concepts in antibody phage display. Briefings in Functional Genomics 2002, 1 (2), 189–203.

50. Yin, Z., Wu, X., Kaczanowska, K., Sungsuwan, S., Comellas Aragones, M., Pett, C., Yu, J., Baniel, C., Westerlind, U., Finn, M. G., Huang, X., Antitumor Humoral and T Cell Responses by Mucin-1 Conjugates of Bacteriophage Qbeta in Wild-type Mice. ACS Chem. Biol. 2018, 13 (6), 1668–1676.

51. Brune, K. D., Leneghan, D. B., Brian, I. J., Ishizuka, A. S., Bachmann, M. F., Draper, S. J., Biswas, S., Howarth, M., Plug-and-Display: decoration of Virus-Like Particles via isopeptide bonds for modular immunization. Sci. Rep. 2016, 6.

52. Bruun, T. U. J., Andersson, A. C., Draper, S. J., Howarth, M., Engineering a Rugged Nanoscaffold To Enhance Plug-and-Display Vaccination. ACS Nano 2018, 12 (9), 8855–8866.

53. Tan, T. K., Rijal, P., Rahikainen, R., Keeble, A., Schimanski, L., Hussain, S., Harvey, R., Hayes, J., Edwards, J., McLean, R., A COVID-19 vaccine candidate using SpyCatcher multimerization of the SARS-CoV-2 spike protein receptor-binding domain induces potent neutralising antibody responses. bioRxiv 2020.

54. Minoshima, F., Ozaki, H., Tateno, H., Integrated analysis of glycan and RNA in single cells. bioRxiv 2020, 2020.06.15.153536.

55. Sterner, E., Flanagan, N., Gildersleeve, J. C., Perspectives on Anti-Glycan Antibodies Gleaned from Development of a Community Resource Database. ACS Chem. Biol. 2016, 11 (7), 1773–83.

56. Gestwicki, J. E., Cairo, C. W., Strong, L. E., Oetjen, K. A., Kiessling, L. L., Influencing receptor-ligand binding mechanisms with multivalent ligand architecture. Journal of the American Chemical Society 2002, 124 (50), 14922–14933.

57. Demetriou, M., Granovsky, M., Quaggin, S., Dennis, J. W., Negative regulation of T-cell activation and autoimmunity by Mgat5 N-glycosylation. Nature 2001, 409 (6821), 733–9.

58. Cecioni, S., Imberty, A., Vidal, S., Glycomimetics versus multivalent glycoconjugates for the design of high affinity lectin ligands. Chem Rev 2015, 115 (1), 525–61.

59. Park, S., Gildersleeve, J. C., Blixt, O., Shin, I., Carbohydrate microarrays. Chem. Soc. Rev. 2013, 42 (10), 4310–4326.

60. Rillahan, C. D., Paulson, J. C., Glycan Microarrays for Decoding the Glycome. Annual Review of Biochemistry 2011, 80 (1), 797–823.

61. Godula, K., Rabuka, D., Nam, K. T., Bertozzi, C. R., Synthesis and microcontact printing of dual end-functionalized mucin-like glycopolymers for microarray applications. Angew Chem Int Ed Engl 2009, 48 (27), 4973–6.

62. Godula, K., Bertozzi, C. R., Density variant glycan microarray for evaluating cross-linking of mucin-like glycoconjugates by lectins. J. Am. Chem. Soc. 2012, 134 (38), 15732–42.

63. Oyelaran, O., Li, Q., Farnsworth, D., Gildersleeve, J. C., Microarrays with Varying Carbohydrate Density Reveal Distinct Subpopulations of Serum Antibodies. J. Proteome Res. 2009, 8 (7), 3529–3538.

64. Yip, Y. L., Hawkins, N. J., Smith, G., Ward, R. L., Biodistribution of filamentous phage-Fab in nude mice. J. Immunol. Methods 1999, 225 (1-2), 171–8.

65. Alam, M. M., Jarvis, C. M., Hincapie, R., McKay, C. S., Schimer, J., Sanhueza, C. A., Xu, K., Diehl, R. C., Finn, M., Kiessling, L. L., Glycan-Modified Virus-like Particles Evoke T Helper Type 1-like Immune Responses. ACS nano 2020.

