## Supporting Information for "DNA-Encoded Multivalent Display of Protein Tetramers on Phage: Synthesis and *In Vivo* Aplications"

### 1. Material and Methods

**1.1** *SpyCatcher_Erwinase (ScA) Cloning and Expression*

SpyCatcher gene (GenBank JQ478411) (Supplementary Table S1d) was synthetized by Integrated DNA Technologies, amplified by PCR using primers SpyCatcher_FW and SpyCatcher_RV (Supplementary Table S1c) and cloned in Erwinase_pET15b vector by restriction enzyme digest and ligation to produce vector containing ScA gene. Vector was transformed by heat shock in chemically competent *E. coli* DH5α cells for amplification and later in chemically competent *E. coli* BL21 (DE3) cells for protein expression.

For protein expression, overnight *E. coli* BL21 (DE3) 20 mL cell culture with 50 µg/mL of carbenicillin was diluted 10-fold into 200 mL of same LB medium and incubated for 1 hour at 37 ºC at 200 rpm. Expression was then induced with a final concentration of 1 mM of IPTG for 3 hours at 200 rpm.

**1.2** *Erwinase and ScA Purification*

*E. coli* BL21(DE3) cells were harvested at 3,220 *g* for 20 min at 4° C. Pellet was resuspended in binding buffer (20 mM $K_3PO_4$, 20 mM imidazole and 500 mM NaCl, pH 7.4) and sonicated for 10 min using constant pulse ON for 15 seg and pulse OFF for 45 seg.

Cell lysate was harvested at 4,000 *g* for 20 min and supernatant was submitted to immobilized metal affinity chromatography. Sample was manually loaded on a 5 mL HisTrap HP His tag protein purification column (GE) and non-specific bound proteins were washed using washing buffer (20 mM potassium phosphate, 200 mM imidazole and 500 mM NaCl, pH 7.4). Either Erwinase or ScA bound proteins were eluted using elution buffer solution (20 mM potassium phosphate, 500 mM imidazole and 500 mM NaCl, pH 7.4).

**1.3** *Construction of SpyTag Phage Vectors*

Degenerated SpyTag sequence oligos including either *BsaI* or *SfiI* restriction enzyme sites (for p3-st and p8-st phages, respectively) were purchased from Integrated DNA Technologies (Supplementary Table S1a). 200 pmol of each library was annealed with 3 molar equivalents of either p3-st or p8-st primers (Supplementary Table S1c). 25 µM of the resulting annealed duplex was extended using 15 U of Klenow fragment, 0.4 mM dNTPs and 1X NEBuffer™ 2 for 20 min at 37 ºC and heat inactivated for 20 min at 70 ºC. Extended double-stranded DNA was purified with Monarck® PCR and DNA Cleanup Kit (New England BioLabs®).

1 µg of purified double-stranded DNA library for p3-st phage vector construction as well as modified M13KE vector were each digested with 20 U of *BsaI-HFv2* and 1X CutSmart© Buffer for 15 min at 37 ºC and heat inactivated for 20 min at 80 ºC. 20 ng of digested insert was ligated with 100 ng of digested vector (24:1 molar ratio) by 5 Weiss U of T4 DNA ligase (Thermo Scientific) in 1X T4 DNA ligase buffer at room temperature overnight.

In addition, 1 µg of purified double-stranded DNA library for p8-st vector construction as well as 450 ng of fth1 vector were each digested with 20 U of *SfiI* and 1X CutSmart© Buffer for 60 min at 50 ºC. Library duplex were purified using Monarck® PCR and DNA Cleanup Kit (New England BioLabs®) and digested vector were loaded on a 0.6% agarose gel and purified using Monarch® DNA Gel Extraction Kit (New England BioLabs®). 1.5 ng of digested insert was ligated with 50 ng of digested vector (4:1 molar ratio) by 5 Weiss U of T4 DNA ligase (Thermo Scientific) in 1X T4 DNA ligase buffer at room temperature overnight.

**1.4** *Isolation of p3-st Phage Clones*

*E. coli* 10G electrocompetent cells were electroporated with 2 µL of ligation product using GenePulser XcellTM Electroporation System. Cells were rapidly resuspended in 500 µL of SOC medium and incubated in a shaker for 45 min at 200 rpm at 37 ºC. Cells were diluted 10-fold in LB medium and 10 µL of the resulting sample was mixed with 200 µL of overnight culture of *E. coli* ER2738 cells. Mixture was plated on LB top agar plates overnight at 37 ºC. Randomly chosen individual plaques formed in the following day were amplified and single-stranded phage DNA was purified and submitted for sequencing.

**1.5** *Isolation of a p8-st Phage Clone*

*E. coli* 10G electrocompetent cells were electroporated with 2 µL of ligation product using GenePulser XcellTM Electroporation System. Cells were rapidly resuspended in 500 µL of SOC medium and incubated in a shaker for 45 min at 200 rpm at 37 ºC. Cells were plated on a tetracycline plate and then incubated overnight at 37 °C. Individual bacterial colonies formed in the following day were inoculated in LB medium and DNA was miniprepped and later submitted for sequencing.

**1.6** *Phage Amplification*

Overnight *E. coli* ER2738 cell culture was used for phage amplification. Phage clone from different sources (plaques, glycerol phage stocks or transformed 10G cells) was mixed with 25 mL of the 100-fold diluted overnight bacteria culture grown to log phage for 1 hour and incubated at 200 rpm at 37 ºC for 5 hours. Culture was centrifuged twice at 4,000 *g* for 15 min and 5 mL of 30% PEG 8,000 / 3M NaCl solution was added to the supernatant. Sample was placed in 4 ºC fridge overnight. In the next day, phage precipitate was collected by centrifuging the solution at 14,000 *g* for 30 min. Pellet was dissolved in 1 mL PBS medium and centrifuged again at 21,000 *g* for 20 min to ensure no cell debris remained in the solution. Supernatant was then transferred to a clean eppendorf tube and placed into a 55 ºC heat block for 10 min. 110 µL of a 10-fold diluted triton X100 solution was added and the solution was then incubated for 1 hour at room temperature. 1/6 volume of 30% PEG 8,000 / 3M NaCl solution was added and precipitate was incubated on ice for 1 hour. Finally, solution was centrifuged at 14,000 *g* for 20 min and remaining pellet was dissolved in either PBS or 1:1 PBS/Glycerol solution for -20 ºC stock.

Expression of recombinant p8 protein fused to SpyTag was analyzed by MALDI-TOF, using a Voyager Elite MALDI mass spectrometer (AB Sciex, US).

**1.7** *Purification of Single-Stranded Phage DNA*

Phage clone was inoculated in 1 mL of 100-fold diluted overnight *E. coli* ER2738 cell culture and incubated at 37 ºC for 5 hours at 200 rpm. Culture was then centrifuged at 2,236 *g* for 15 min at room temperature and supernatant was transferred to a clean eppendorf tube. Centrifugation was repeated to ensure removal of all bacterial cells. Isolation of single-stranded DNA was then performed following QIAprep® Spin M13 Kit instructions.

**1.8** *Conjugation of Erwinase to Phages using SpyTag/SpyCatcher System*

All conjugations were performed at room temperature for 1 hour. All phage and protein solutions were transferred to PBS buffer and mixed together at desired proportions. Protein and phage concentrations were estimated spectrophotometrically at 280nm and 269/320 nm, respectively. Bacterial lysates in binding buffer (20 mM potassium phosphate, 20 mM imidazole and 500 mM NaCl, pH 7.4) were transferred to PBS buffer using ZebaTM Spin Desalting column (Thermo Scientific).

**1.9** *In Vivo Studies and Blood Sample Processing*

All the procedures and experiments involving animals were carried out using a protocol approved by the Health Sciences Laboratory Animal Services (HSLAS), University of Alberta. The protocol was approved as per the Canadian Council on Animal Care (CCAC) guidelines. All mice were maintained in pathogen-free conditions at the University of Alberta breeding facility. Animals were injected with protein-phage conjugates (0.1 mL, $10^{12}$ PFU/mL (first *in vivo* experiment) or $10^{11}$ PFU/mL – second *in vivo* experiment – in PBS). 50 µL of blood sample was drawn at different time points from each mouse. Two hours post-injection mice were euthanized. Blood samples were centrifuged for 5 min at 2,000 *g*. Plasma was collected to determine the amount of phage by sequencing, plaque forming unit (PFU) and/or colony forming unit (CFU) assays.

**1.10** *qPCR*

Phage samples were mixed in reaction mix containing home-made Taq DNA polymerase, 0.2 mM dNTPs, 3X SBRY1, 5% v/v DMSO, 0.2 µM NF10 forward primer and 0.2 µM '96' reverse primer (for sequence, see Supplementary Table S1c) in 1X Phusion® HF Reaction Buffer (Thermo Scientific). qPCR reaction was performed on a Bio-Rad C1000 thermocycler using the following parameters: first step at 95 ºC for 3 min, second step repeated 35 times at 95 ºC for 10 seg, 58 ºC for 30 seg, 72 ºC for 20 seg and 80 ºC for 20 seg. A final melting point step from 65 to 95 ºC was also performed. Data was analyzed using CFX ManagerTM software (Bio-Rad).

**1.11** *Illumina Sequencing*

qPCR amplicons were converted to Illumina-compatible DNA by PCR as described in our previous publications[1, 2]. Each qPCR template was combined with 1X Phusion® HF Reaction Buffer (Thermo Scientific), home-made Taq DNA polymerase, 0.2 mM dNTPs, 5% v/v DMSO,

0.2 µM Illumina forward primer and 0.2 µM Illumina reverse primer (Supplementary Table S1c). Each primer has four variable nucleotide sequences (NNNN) which were used to track multiple Illumina samples. PCR was performed on an Eppendorf Mastercycler thermocycler using the following parameters: first cycle at 95 ºC for 3 min, second step repeated 10 times at 95 ºC for 10 seg, 58 ºC for 30 seg and 72 ºC for 20 seg and a final step at 72 ºC for 20 seg. PCR products were quantified based on intensity from agarose gel, purified and submitted for deep sequencing.

**1.12** *Electrophoresis*

DNA samples were analyzed on 1% agarose gel (1% w/v agarose, 1X SYBR™ Safe DNA Gel Stain (Thermo Scientific), and 1X TAE buffer). Primers and degenerated oligos were analyzed using the same procedure but with agarose concentration increased to 4%. TAE buffer (40 mM Tris, 0.11% acetic acid and 1 mM EDTA, pH 8.0) was used to run samples at 110 V.

Protein expression and conjugation experiments were analyzed on 10% Bis-Tris SDS-PAGE gel with MES running buffer. Resolving gel was made using 10% acrylamide, 0.27% bis-acrylamide, 0.36 M bis-tris pH 6.6, 0.06% w/v APS and 0.0025% v/v TEMED. Stacking gel was prepared with 4.98% acrylamide, 0.13% bis-acrylamide, 0.36 M bis-tris pH 6.6, 0.08% w/v APS and 0.004% v/v TEMED. For MES running buffer, 50 mM MES, 50 mM Tris, 1 mM EDTA, 0.1% w/v SDS and 5 mM sodium bisulfite were used. Samples were run at 35 mA. Gels were stained with 0.3% Coomassie R-250, 10% methanol and 10% acetic acid and distained with 40% methanol / 10% acetic acid solution. Band intensities were calculated using the ImageJ program (National Institutes of Health).

**1.13** *Enzymatic Assay*

All Erwinase enzymatic activities were measuring using Nessler's method. 10 µL of protein sample was incubated with pre-warmed 23.8 mM Tris HCl, pH 8.6, and 9 mM L-asparagine solution at 37 ºC for 10 min. Reaction was stopped with 70 mM trichloroacetic acid (TCA) and diluted 21.5-fold in water. Nessler's reagent (Sigma) was added at a final concentration of 10% v/v and absorbance was measured at 436 nm after 1 min. Enzymatic activity was measured as U/mL where U is defined as the amount of enzyme required to liberate 1 µmol of ammonia from L-asparagine per minute at pH 8.6 at 37 ºC. $(NH_4)_2SO_4$ standard curve was used to quantify activity (Supplementary Figure S8).

**1.14** *Phage Titering*

Phage concentration was estimated by titering. *E. coli* ER2738 cells were grown on LB for 6 hours to mid-log phase. Serial phage dilutions in LB were prepared from different sources (e.g., phage stocks in glycerol, phage in PBS buffer or phage collected from mice plasma) and used for titering. 10 µL of each dilution was added to 200 µL of ER2738 cells and incubated for about 5 min. Infected cells were transferred to melted top agar, vortexed briefly, poured onto LB/IPTG/Xgal plates and incubated overnight at 37 ºC.

Titering of phage was also performed by counting the number of infected TG1 colonies. TG1 cells were firstly grown to log phase. Then, 196 µL of the cells were mixed with 4 µL of 100-fold diluted plasma samples and overgrown for 30 min at 37 °C. From this mixture, 50 µL of cells were plated on tetracycline added LB plates and incubated overnight at 37 °C.

**1.15** *Purification of Protein-Phage Conjugates from Non-Conjugated Proteins*

Purification of protein-phage conjugates from conjugation reaction was performed by simply adding 1/6 volume of 30% PEG 8,000 / 3M NaCl solution to the reaction mix. After 1 hour on ice, solution was centrifuged at 14,000 *g* for 20 min and supernatant was removed. Non-conjugated protein-free pellet was then dissolved in either PBS buffer or 1:1 PBS/Glycerol for -20 ºC long-term storage.

**a**

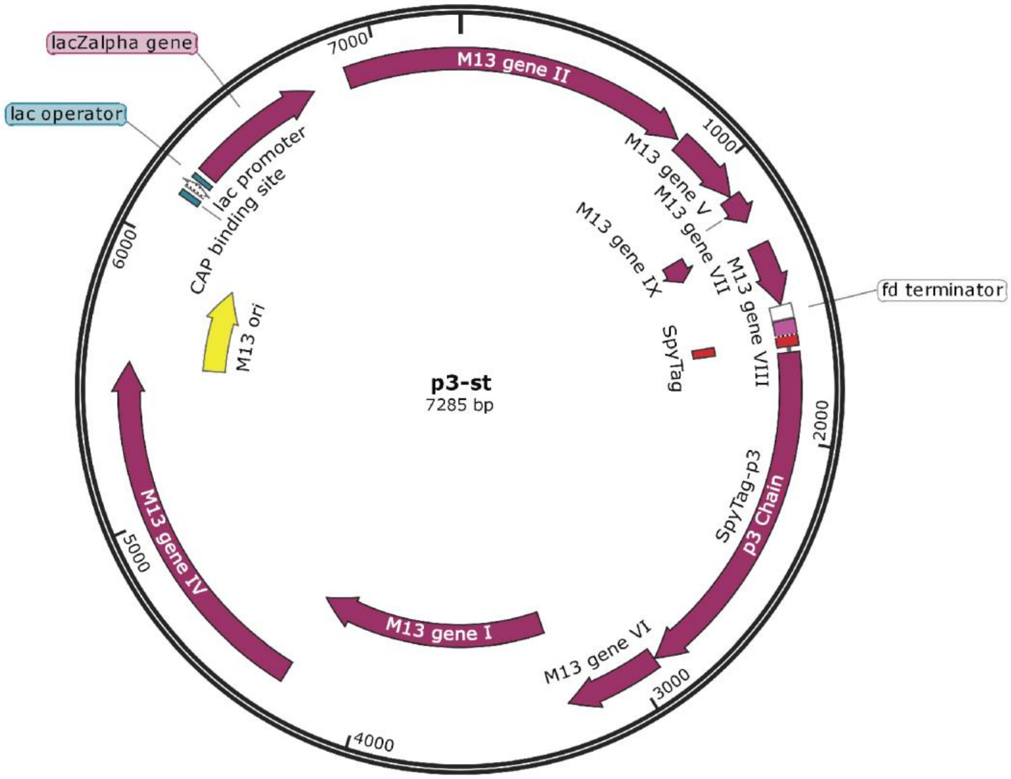

| | | | | | | | | | | | | | |
|---|---|---|---|---|---|---|---|---|---|---|---|---|---|
| p3-st | GCN | CAY | ATH | GTN | ATG | GTN | GAY | GCN | TAY | AAR | CCN | ACN | AAR |
| Clone 1 | GCC | CAC | ATC | GTT | ATG | GTG | GAC | GCA | TAC | AAA | CCT | ACA | AAA |
| Clone 2 | GCC | CAT | ATA | GTT | ATG | GTA | GAC | GCG | TAT | AAA | CCG | ACG | AAA |
| Clone 3 | GCC | CAT | ATA | GTA | ATG | GTT | GAT | GCA | TAC | AAA | CCT | ACG | AAA |

**b**

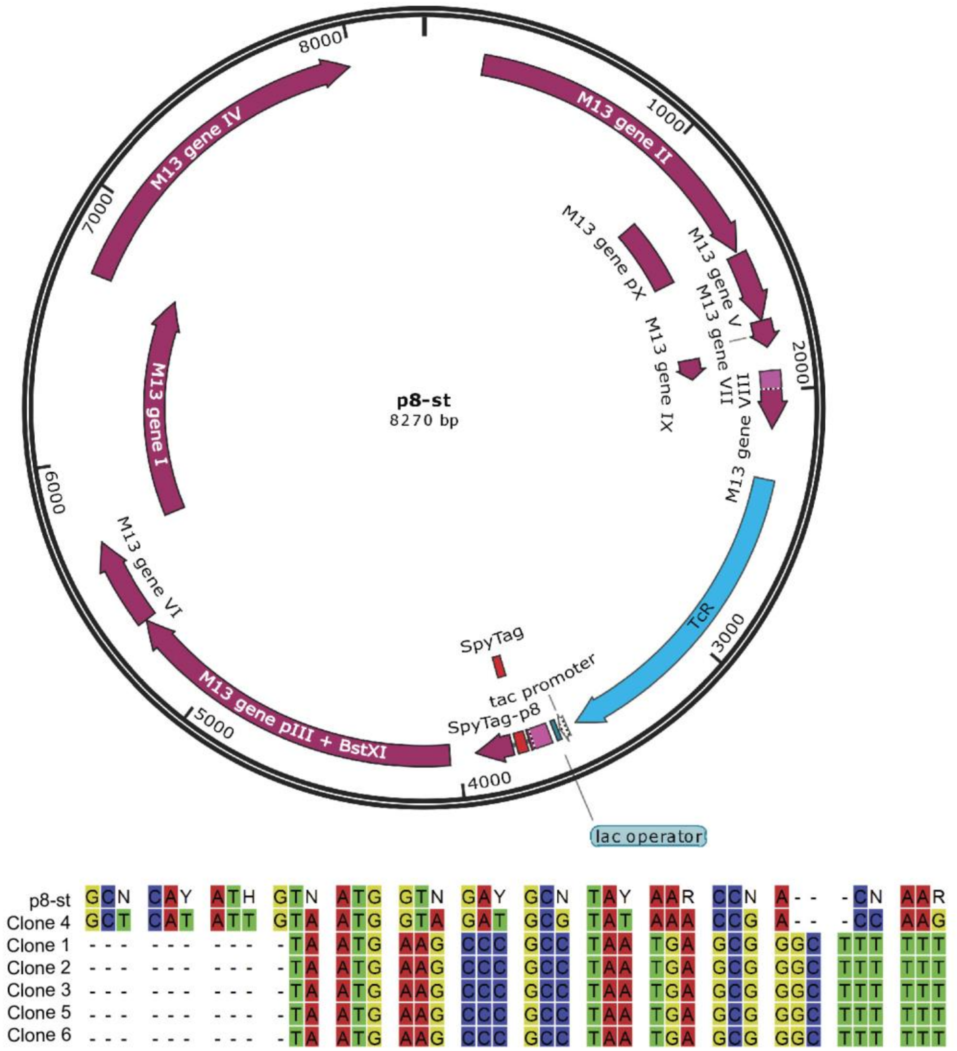

**Supplementary Figure S1.** p3-st (from M13KE) and p8-st (from fth1) bacteriophage vectors. (a) p3-st vector (top) and genetic sequence of degenerated SpyTag and screened p3-st phage clones (bottom). SpyTag sequence was cloned in the N-terminus of the M13 p3 chain gene, right after the M13 p3 signal sequence. (b) p8-st vector (top) and genetic sequence of degenerated SpyTag and screened p8-st bacterial clones (bottom). SpyTag sequence was cloned in the N-terminus of the recombinant fd p8 chain gene, right after the fd p8 signal sequence. Clones 1-3, 5-6 contain no insert. For reference, N = any nucleotide, Y = C or T, H = A or C or T and R = A or G. Plasmid maps were drawn using *SnapGene Viewer*.

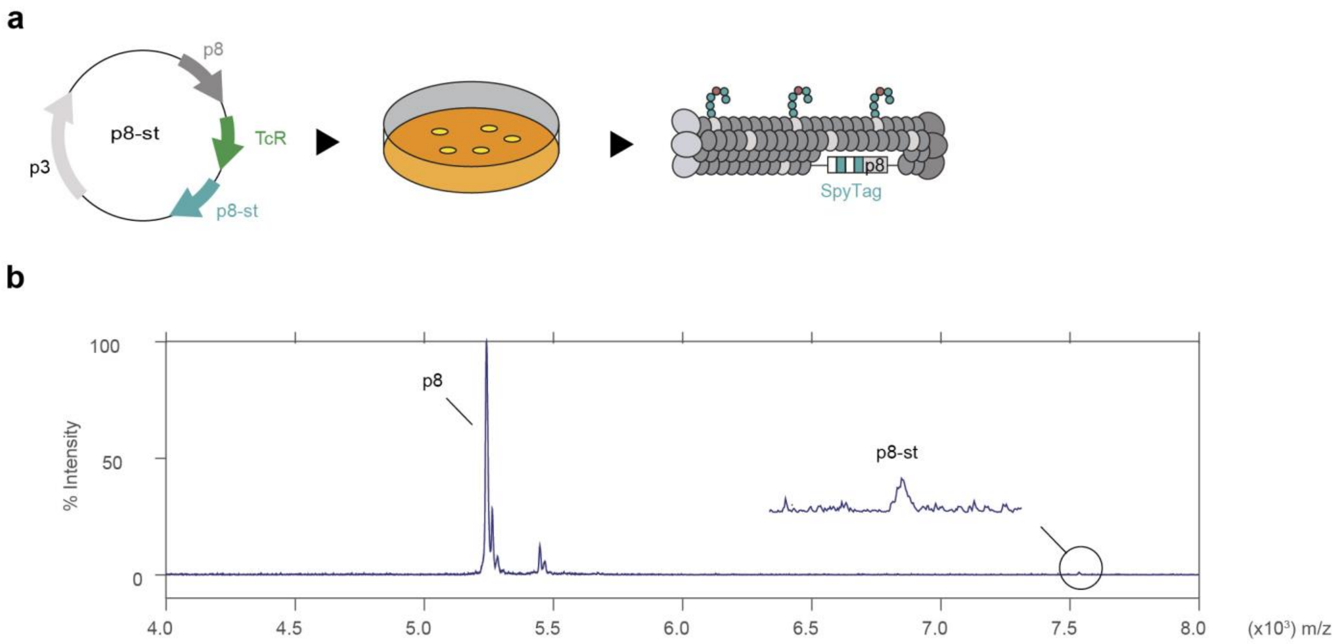

**Supplementary Figure S2.** Design of p8-st phage.
(a) Cloning and isolation of st-p8 phages. TcR = tetracycline resistant gene; TcR marker was used to screen tetracycline resistant bacteria containing the p8-st vector (b) MALDI spectrum of p8 and recombinant p8. Recombinant p8 protein contains the SpyTag sequence fused to its N-terminus position. Relative peak intensity suggests a ratio of approximately 50 copies of SpyTag-displaying recombinant p8 proteins and 2700 copies of the wild-type p8 protein (total number of p8) or 50 copies per phage particle.

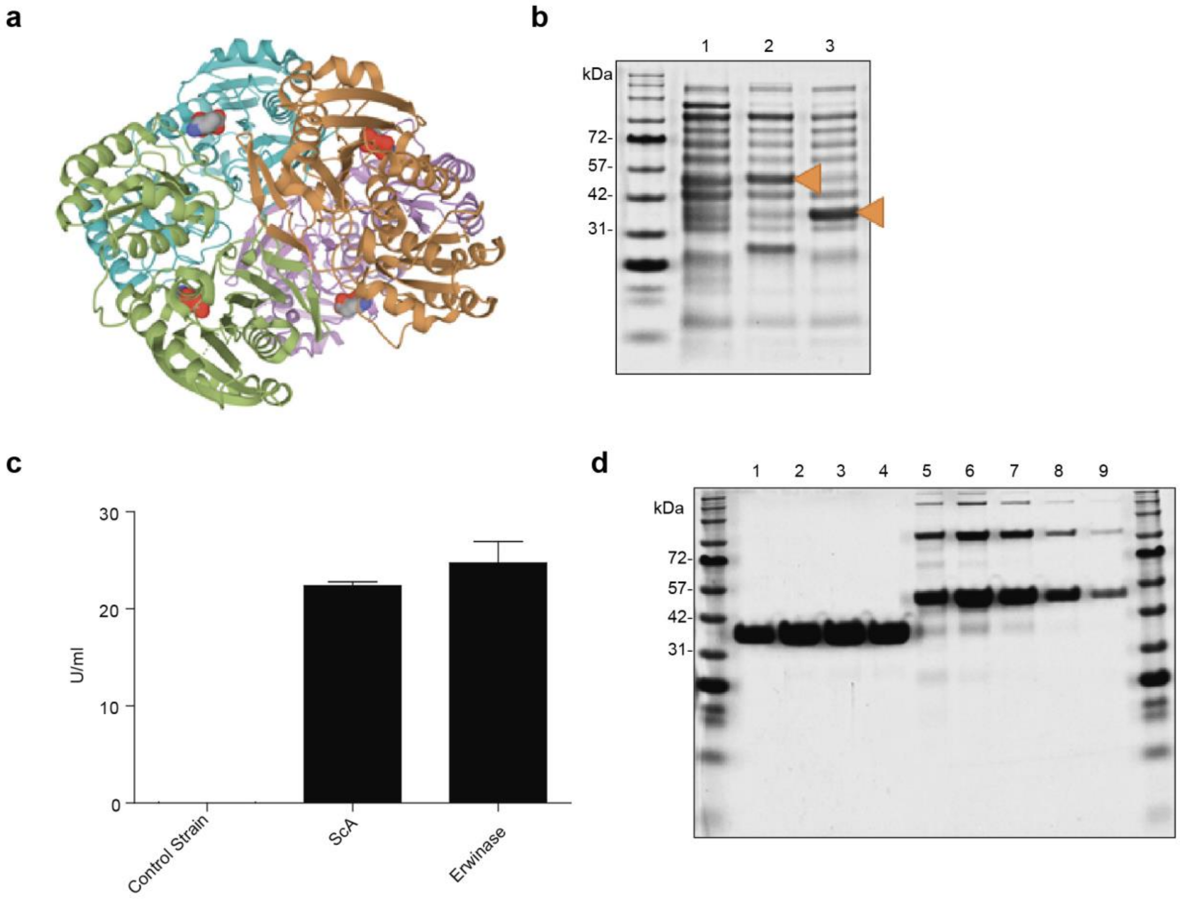

**Supplementary Figure S3.** Expression of ScA.

We have previously shown that fusing a histidine tag to the N-terminus of Erwinase did not significantly affect its enzyme activity[3]. Therefore, we postulated that adding SpyCatcher to the same position would likely not affect the biological activity of our protein, which has been confirmed by this set of experiments. (a) X-ray structure of tetrameric protein Erwinase (cartoon representation coloured by chain) bound to its substrates (surface representation). (b) SDS-PAGE image of soluble fractions originated from BL21(DE3) lysates: (1) BL21(DE3) cells with no plasmid; (2) ScA-expressing BL21(DE3) cells; (3) Erwinase-expressing BL21(DE3) cells. Orange arrows indicate recombinant ScA (left) and non-fused recombinant Erwinase (right) proteins. (c) Enzymatic activity of soluble fraction from BL21(DE3) lysate. ScA and Erwinase enzymatic activities are statistically not different ($p<0.05$). n=3, unpaired t-test (d) Eluted fractions obtained after purification of both recombinant Erwinases. Lanes 1 – 4: Eluted fractions of non-fused recombinant Erwinase. Lanes 5 – 9: Eluted fractions from fusion ScA. We observed ~30% of the ScA protein eluting as a covalent dimer at ~100 kDa. We were not able to remove these bands after size-exclusion chromatography or SDS-PAGE sample treatment with high concentrations of chaotropic salts/DTT/β-mercaptoethanol and increases in denaturing time (Results not shown).

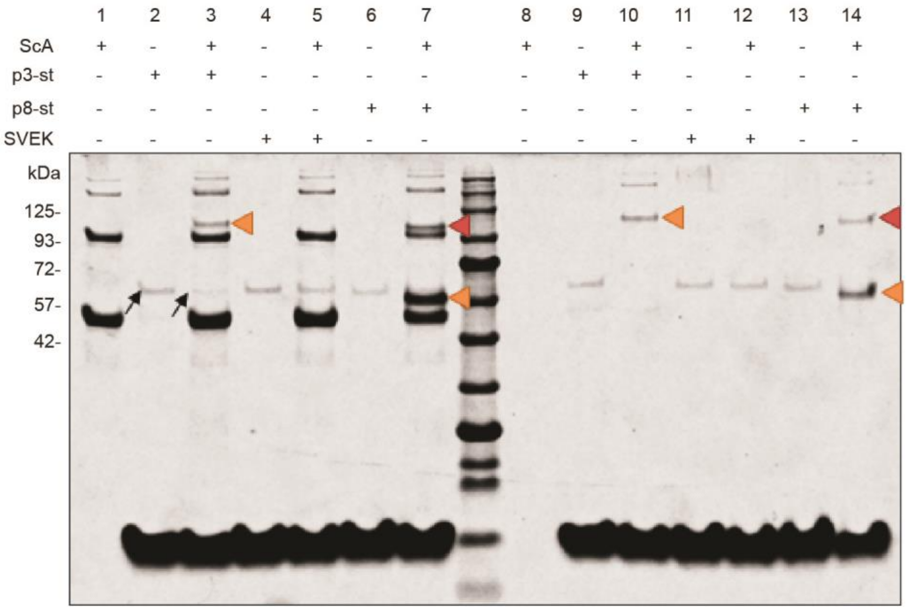

**Supplementary Figure S4.** Conjugation of ScA to SpyTag-Expressing Phages.
SDS-PAGE image representing conjugation reactions before (left-side of molecular weight marker, lanes 1 - 7) and after purification using PEG/NaCl precipitation (right-side of molecular weight marker, lanes 8 - 14). Orange arrows indicate conjugates observed only when mixing ScA with either p3-st or p8-st. Red arrows indicate dimer-phage conjugates. Black arrows indicate consumption of p3 protein from p3-st phage.

**a**

ScA and SpyTag

```
19 SHMAMVD     25
   :|:.|||
 1 AHIVMVD      7
```

**b**

Erwinase and SpyTag

```
116 VVFVAAMRP    124
    :|.|.|.:|
  3 IVMVDAYKP     11
```

**Supplementary Figure S5.** Local alignments between (a) ScA and SpyTag and (b) Erwinase and SpyTag sequences.

We suspected that both extra upper bands observed in ScA eluted fractions (Supplementary Figure S3) could be caused by oligomerization between ScA monomers through ligation of its SpyCatcher portion to a SpyTag-like portion of another monomer. In fact, both band sizes correspond to ScA dimer and tetramer molecular weights (101.2 and 202.4 kDa, respectively). Local alignment between ScA sequence and SpyTag revealed a 57.1% identity and 85.7% similarity between ScA residues 19 and 25 (SHMAMVD) and SpyTag sequence 1 to 7 (AHIVMVD). On the other hand, local alignment of only the Erwinase domain and the SpyTag sequence showed only a 44.4% identity and 66.7% similarity between the Erwinase residues 116 and 124 (VVFVAAMRP) and the SpyTag sequence 3 to 11 (IVMVDAYKP). However, differently from the first alignment, the SpyTag reactive aspartate residue (D) was not present in the Erwinase sequence match, thus indicating that self-reaction between the SpyCatcher domains might be the truly mechanism playing a role in the formation of the observed dimers/tetramers. Therefore, we concluded that the extra bands observed in SDS-PAGE gels are not contaminants but rather covalently bound ScA oligomeric structures, in which SpyCatcher domains from different proteins have self-reacted. This SpyCatcher self-reactivity property has already been observed in previous works involving improved variants of SpyCatcher.[4]

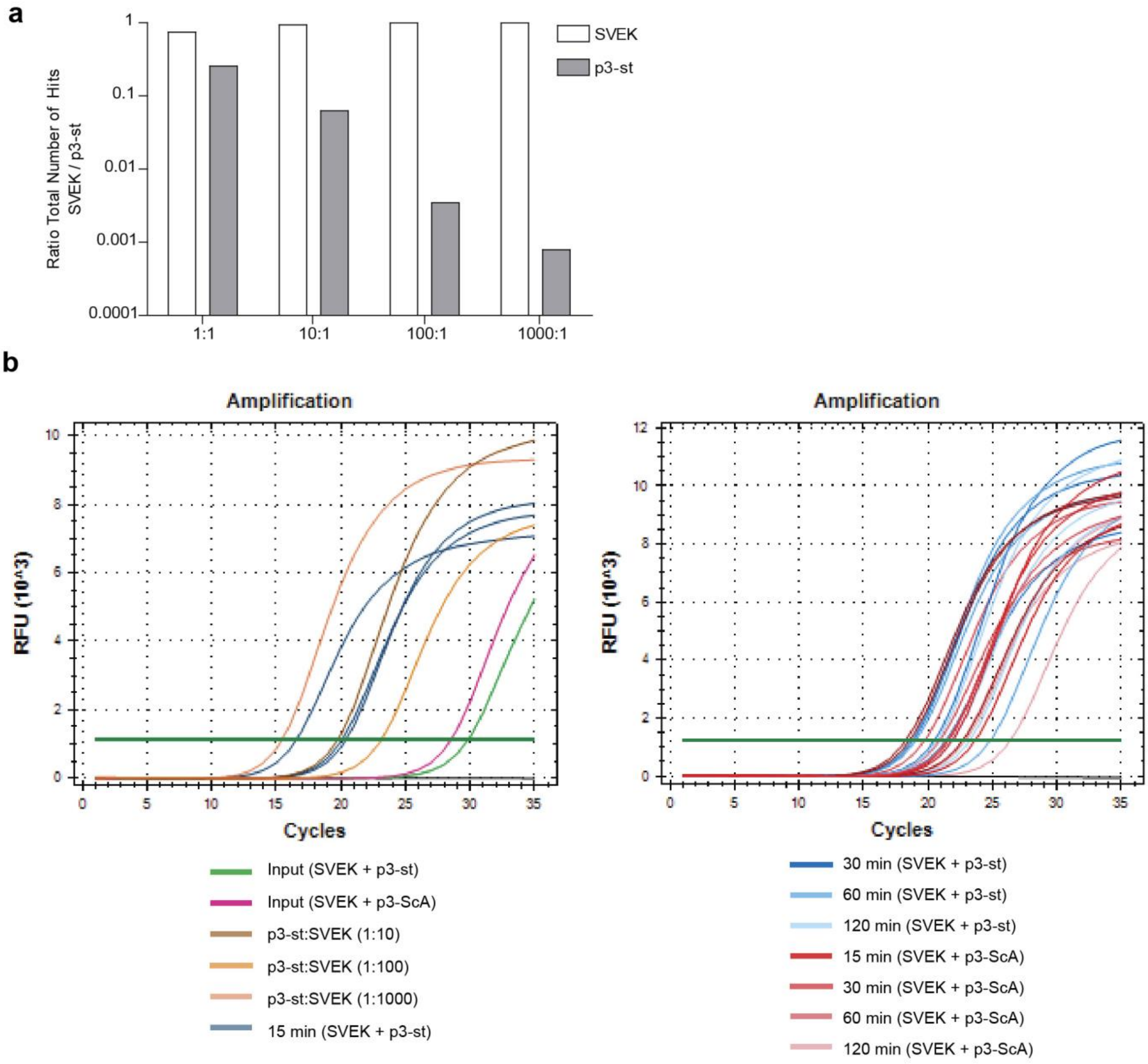

**Supplementary Figure S6.** Amplification and sequencing of input and plasma samples containing SVEK and p3-st/p3-ScA$_5$ phages.

(a) Ratio of number of hits of p3-st and SVEK at different concentrations of each clone detected by Illumina sequencing. (b) qPCR of input and plasma samples. Input are samples used for injection containing the described phages. Also, individual phage clones (SVEK and p3-st) were mixed at different ratios. Blue and red curves represent qPCR of diluted plasma samples at different time points (15, 30 60 and 120 min) for mice treated with protein-phage conjugates or phage alone, respectively. (c) Clearance of phages from plasma as measured by the number of hits from each clone collected from deep sequencing data. Repeated Measures ANOVA statistical was used to test for significance.

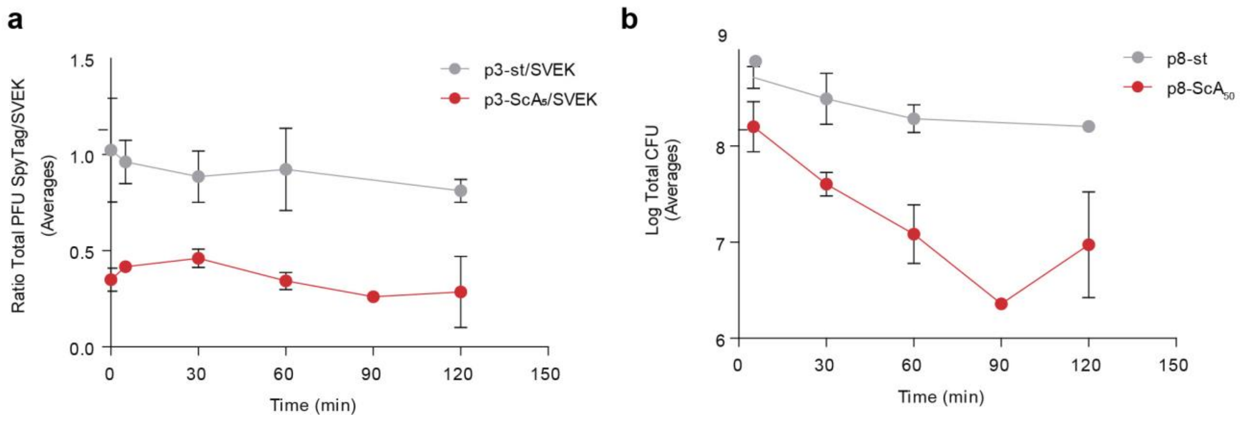

**Supplementary Figure S7.** Clearance of protein-phage conjugates as measured by average of ratio of total PFUs or total CFUs.

**(a)** Clearance of p3-st/p3-Sc$A_5$ phages from plasma as measured by ratio of number of plaques from each clone (p3-st/SVEK or p3-Sc$A_5$/SVEK). Repeated Measures ANOVA statistical analysis was used to test for significance from time = 0 (initial sample) to time = 60 min. **(b)** Clearance of p8-st/p8-Sc$A_{50}$ from plasma. Repeated Measures ANOVA statistical analysis was used to test for significance from time = 5 min to time = 60 min. Measurements represent means ± SD.

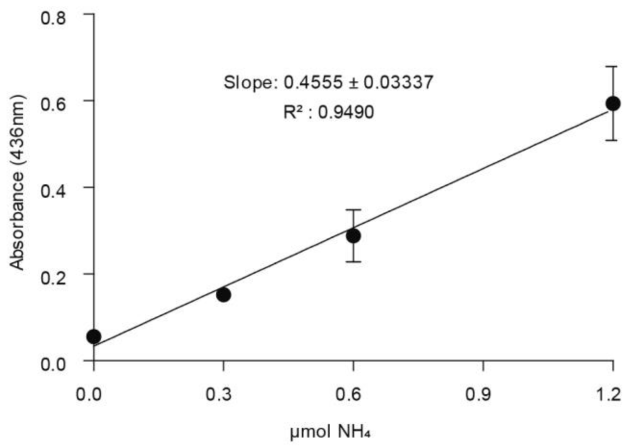

**Supplementary Figure S8.** Nessler's assay calibration curve. Ammonium sulfate was used as a standard.

**Supplementary Table S1.** Sequence of degenerated oligos, protein primary structures, PCR primers, and genes.

a) Degenerated Oligos

| Name | Sequence |
|---|---|
| Degenerated Oligo for p3-st vector | GCGCCCGGCCGAGGTCTCGCCACCACTTTCACCACTACCYTTNGTNGGYTTRTANGCRTCNACCATNACDATRTGNGCAGAGTAGAGACCGTGAG |
| Degenerated Oligo for p8-st vector | GTTTCTTGGCCAACGTGGCCGGGCNCAYATHGTNATGGTNGAYGCNTAYAARCCNACNAARTTGGCCTCTGGGGCCAAGAAAC |

b) Protein Primary Structures

| Name | Sequence |
|---|---|
| SpyCatcher | AMVDTLSGLSSEQGQSGDMTIEEDSATHIKFSKRDEDGKELAGATMELRDSSGKTISTWISDGQVKDFYLYPGKYTFVETAAPDGYEVATAITFTVNEQGQVTVNGKATKGDAHI |
| ScA | MGSSHHHHHHSSGLVPRGSHMAMVDTLSGLSSEQGQSGDMTIEEDSATHIKFSKRDEDGKELAGATMELRDSSGKTISTWISDGQVKDFYLYPGKYTFVETAAPDGYEVATAITFTVNEQGQVTVNGKATKGDAHIGGGGSGGGGSHMADKLPNIVILATGGTIAGSAATGTQTTGYKAGALGVDTLINAVPEVKKLANVKGEQFSNMASENMTGDVVLKLSQRVNELLARDDVDGVVITHGTDTVEESAYFLHLTVKSDKPVVFVAAMRPATAISADGPMNLLEAVRVAGDKQSRGRGVMVVLNDRIGSARYITKTNASTLDTFKANEEGYLGVIIGNRIYYQNRIDKLHTTRSVFDVRGLTSLPKVDILYGYQDDPEYLYDAAIQHGVKGIVYAGMGAGSVSVRGIAGMRKAMEKGVVVIRSTRTGNGIVPPDEELPGLVSDSLNPAHARILLMLALTRTSDPKVIQEYFHTY |
| p3 fused to SpyTag | AHIVMVDAYKPTKGSGESGGSAETVESCLAKSHTENSFTNVWKDDKTLDRYANYEGCLWNATGVVVCTGDETQCYGTWVPIGLAIPENEGGGSEGGGSEGGGSEGGGTKPPEYGDTPIPGYTYINPLDGTYPPGTEQNPANPNPSLEESQPLNTFMFQNNRFRNRQGALTVYTGTVTQGTDPVKTYYQYTPVSSKAMYDAYWNGKFRDCAFHSGFNEDLFVCEYQGQSSDLPQPPVNAGGGSGGGSGGGSEGGGSEGGGSEGGGSEGGGSGGGSGSGDFDYEKMANANKGAMTENADENALQSDAKGKLDSVATDYGAAIDGFIGDVSGLANGNGATGDFAGSNSQMAQVGDGDNSPLMNNFRQYLPSLPQSVECRPFVFGAGKPYEFSIDCDKINLFRGVFAFLLYVATFMYVFSTFANILRNKES |
| p8 fused to SpyTag | AEGGQRGRAHIVMVDAYKPTKLASGADPAKAAFDSLQASATEYIGYAWAMVVVIVGATIGIKLFKKFTSKAS |
| SVEK | SVEKNDQKTYHAGGG |

### c) PCR Primers

| Name | Sequence |
|---|---|
| SpyCatcher_FW | GTTTCTTCCATATGGCCATGGTTGATACCTTATCAGG |
| SpyCatcher_RV | GTTTCTTCCATATGACTGCCACCGCCACCGCTACCGCCACCGCCAATAT<br>GAGCGTCACCTTTAGTTGC |
| p3-st | CATGCCCGGGTACCTTTCTATTCTCACGGTCTCTACTCTGC |
| p8-st | GTTTCTTGGCCCCAGAGG |
| NF10 | TTTTGGAGATTTTCAACGTG |
| ‘96’ | CCCTCATAGTTAGCGTAACG |
| Illumina Forward | CAAGCAGAAGACGGCATACGAGATCGGTCTCGGCATTCCTGCTGAACC<br>GCTCTTCCGATCTNNNNTTGGAGATTTTCAACGTG |
| Illumina Reverse | AATGATACGGCGACCACCGAGATCTACACTCTTTCCCTACACGACGCTC<br>TTCCGATCTNNNNACAGTTTCGGCCGA |

### d) Genes

| Name | Sequence |
|---|---|
| SpyCatcher | GCCATGGTTGATACCTTATCAGGTTTATCAAGTGAGCAAGGTCAGTCCGGTGAT<br>ATGACAATTGAAGAAGATAGTGCTACCCATATTAAATTCTCAAAACGTGATGAG<br>GACGGCAAAGAGTTAGCTGGTGCAACTATGGAGTTGCGTGATTCATCTGGTAAA<br>ACTATTAGTACATGGATTTCAGATGGACAAGTGAAAGATTTCTACCTGTATCCA<br>GGAAAATATACATTTGTCGAAACCGCAGCACCAGACGGTTATGAGGTAGCAACT<br>GCTATTACCTTTACAGTTAATGAGCAAGGTCAGGTTACTGTAAATGGCAAAGCA<br>ACTAAAGGTGACGCTCATATT |
| ScA | ATGGGCAGCAGCCATCATCATCATCATCACAGCAGCGGCCTGGTGCCGCGCGGC<br>AGCCATATGGCCATGGTTGATACCTTATCAGGTTTATCAAGTGAGCAAGGTCAG<br>TCCGGTGATATGACAATTGAAGAAGATAGTGCTACCCATATTAAATTCTCAAAA<br>CGTGATGAGGACGGCAAAGAGTTAGCTGGTGCAACTATGGAGTTGCGTGATTCA<br>TCTGGTAAAACTATTAGTACATGGATTTCAGATGGACAAGTGAAAGATTTCTAC<br>CTGTATCCAGGAAAATATACATTTGTCGAAACCGCAGCACCAGACGGTTATGAG<br>GTAGCAACTGCTATTACCTTTACAGTTAATGAGCAAGGTCAGGTTACTGTAAAT<br>GGCAAAGCAACTAAAGGTGACGCTCATATTGGCGGTGGCGGTAGCGGTGGCGG<br>TGGCAGTCATATGGCCGACAAGCTGCCGAATATCGTCATCCTGGCAACCGGCGG<br>TACGATTGCGGGTTCAGCCGCAACCGGTACCCAGACCACCGGTTATAAAGCAGG<br>TGCTCTGGGCGTTGACACCCTGATCAACGCGGTGCCGGAAGTTAAAAAGCTGGC<br>CAATGTGAAAGGCGAACAGTTTTCAAACATGGCCTCGGAAAATATGACCGGTG<br>ATGTGGTTCTGAAGCTGTCCCAACGTGTGAACGAACTGCTGGCCCGCGATGACG<br>TTGATGGCGTCGTGATTACCCATGGTACCGACACGGTTGAAGAAAGTGCATACT<br>TTCTGCACCTGACGGTCAAATCCGATAAGCCGGTTGTCTTCGTGGCAGCGATGC<br>GTCCGGCCACCGCAATCAGCGCCGATGGTCCGATGAACCTGCTGGAAGCCGTCC<br>GTGTGGCAGGTGACAAACAGTCTCGTGGCCGCGGTGTGATGGTGGTTCTGAATG<br>ATCGTATTGGCTCAGCACGCTATATCACCAAAACGAACGCTTCGACCCTGGATA<br>CGTTTAAGGCGAATGAAGAAGGTTACCTGGGCGTTATCATCGGTAACCGTATCT<br>ACTACCAAAACCGCATCGATAAGCTGCATACCACCCGTAGCGTCTTTGATGTGC<br>GCGGCCTGACCTCTCTGCCGAAGGTGGACATTCTGTATGGTTACCAGGATGACC<br>CGGAATATCTGTACGATGCGGCCATTCAACACGGTGTTAAAGGCATCGTCTATG<br>CAGGCATGGGTGCTGGCAGCGTTTCTGTCCGTGGTATCGCGGGCATGCGCAAAG<br>CCATGGAAAAGGGCGTCGTGGTTATTCGTAGCACCCGTACCGGTAATGGTATTG<br>TTCCGCCGGATGAAGAACTGCCGGGTCTGGTCAGCGACTCCCTGAATCCGGCAC<br>ATGCTCGTATTCTGCTGATGCTGGCACTGACCCGCACGTCCGACCCGAAAGTTA<br>TCCAAGAATACTTCCACACCTACTGA |
| SVEK | AGTGTTGAGAAAAATGATCAAAAAACTTATCACGCCGGTGGTGGT |

**Supplementary Table S2.** Number of plaques/colonies observed after titering of plasma samples.

M1 = mouse 1 (control), M2 = mouse 2 (control), M3 = mouse 3 (control), M4 = mouse 1 (treated), M5 = mouse 2 (treated) and M6 = mouse 3 (treated). # = number of plaques (for p3-st, p3-$ScA_5$ and SVEK phages) or number of colonies (for p8-st and p8-$ScA_{50}$ phages). DF = dilution factor. * measurement was taken after 90 min instead of 120 min.

| | 5 min | | 30 min | | 60 min | | 120 min | |
|---|---|---|---|---|---|---|---|---|
| Phage Titered | # | DF | # | DF | # | DF | # | DF |
| p3-st (M1) | 265 | 100000 | 185 | 100000 | 158 | 100000 | | |
| SVEK (M1) | 303 | 100000 | 216 | 100000 | 233 | 100000 | | |
| p8-st (M1) | >5000 | 100 | ~1500 | 100 | ~1500 | 100 | | |
| p3-st (M2) | 357 | 10000 | 80 | 10000 | 121 | 10000 | 80 | 10000 |
| SVEK (M2) | 387 | 10000 | 104 | 10000 | 119 | 10000 | 104 | 10000 |
| p8-st (M2) | ~3200 | 100 | >5000 | 100 | ~2300 | 100 | ~1330 | 100 |
| p3-st (M3) | 49 | 100000 | 34 | 100000 | 44 | 100000 | 35 | 100000 |
| SVEK (M3) | 45 | 100000 | 33 | 100000 | 41 | 100000 | 41 | 100000 |
| p8-st (M3) | >5000 | 100 | ~2300 | 100 | ~1200 | 100 | ~1340 | 100 |
| p3-$ScA_5$ (M4) | 139 | 100000 | 103 | 100000 | 27 | 100000 | 36* | 100000 |
| SVEK + ScA (M4) | 317 | 100000 | 253 | 100000 | 70 | 100000 | 138* | 100000 |
| p8-$ScA_{50}$ (M4) | ~2300 | 100 | 270 | 100 | 68 | 100 | 19* | 100 |
| p3-$ScA_5$ (M5) | 115 | 100000 | 72 | 100000 | 64 | 100000 | 17 | 100000 |
| SVEK + ScA (M5) | 281 | 100000 | 145 | 100000 | 185 | 100000 | 110 | 100000 |
| p8-$ScA_{50}$ (M5) | ~1440 | 100 | 300 | 100 | 67 | 100 | 32 | 100 |
| p3-$ScA_5$ (M6) | 27 | 100000 | 22 | 100000 | 146 | 10000 | 15 | 100000 |
| SVEK + ScA (M6) | 67 | 100000 | 46 | 100000 | 491 | 10000 | 36 | 100000 |
| p8-$ScA_{50}$ (M6) | ~700 | 100 | 460 | 100 | 228 | 100 | 192 | 100 |

### References

1. Sojitra, M.; Sarkar, S.; Maghera, J.; Rodrigues, E.; Carpenter, E.; Seth, S.; Vinals, D. F.; Bennett, N.; Reddy, R.; Khalil, A.; Xue, X.; Bell, M.; Zheng, R. B.; Zhang, P.; Nycholat, C.; Ling, C.-C.; Lowary, T. L.; Paulson, J. C.; Macauley, M. S.; Derda, R., Genetically Encoded, Multivalent Liquid Glycan Array (LiGA). *bioRxiv* **2020**, 2020.03.24.997536.
2. He, B.; Tjhung, K. F.; Bennett, N. J.; Chou, Y.; Rau, A.; Huang, J.; Derda, R., Compositional Bias in Naive and Chemically-modified Phage-Displayed Libraries uncovered by Paired-end Deep Sequencing. *Sci. Rep.* **2018,** *8* (1), 1214.
3. Wlodarczyk, S. R.; Costa-Silva, T. A.; Pessoa-Jr, A.; Madeira, P.; Monteiro, G., Effect of osmolytes on the activity of anti-cancer enzyme L-Asparaginase II from Erwinia chrysanthemi. *Process Biochem.* **2019,** *81*, 123-131.
4. Keeble, A. H.; Banerjee, A.; Ferla, M. P.; Reddington, S. C.; Anuar, I.; Howarth, M., Evolving Accelerated Amidation by SpyTag/SpyCatcher to Analyze Membrane Dynamics. *Angewandte Chemie-International Edition* **2017,** *56* (52), 16521-16525.
